# Profiling the secretome: maternal obesity impacts redox and adipogenic signaling molecules during neonatal mesenchymal stem cells’ adipogenesis

**DOI:** 10.1101/2025.04.21.649774

**Authors:** Sofía Bellalta, Erika Pinheiro-Machado, Paola Casanello, Marijke Faas, Torsten Plösch

## Abstract

Recent studies evidence an altered bioenergetic profile and higher adipogenic commitment in the MSCs from neonates of mothers with obesity. We hypothesize that these alterations may also affect the secretome of these cells. The aim of this study was to characterize the redox and adipogenic secretome of MSCs from the offspring of women with obesity compared to the ones from normal-weight women, both before and during adipogenesis. Wharton’s jelly-derived MSCs were isolated from newborns of normal-weight women (NW-MSC; Body Mass Index 18.5-24.5 kg/m²) and women with obesity (OB-MSC; Body Mass Index>30 kg/m²) and cultured for 0, 5 and 21 days of adipogenesis. The secretome from these cells was collected during the three timepoints and characterized through mass spectrometry. Our findings reveal fundamental difference in the secretome profiles, primarily associated with pathways involved in cellular and metabolic processes (p<0.05). Maternal obesity was found to decrease redox capacity at day 0 but subsequently triggered a compensatory increase in redox proteins during adipogenesis of OB-MSCs (p<0.05). Additionally, OB-MSCs secreted higher levels of lipid synthesis proteins and proinflammatory adipokines (p<0.05), which may contribute to the dysregulated adipogenesis observed in obesity. These results indicate that maternal obesity programs the secretome of neonatal MSCs, supporting that maternal obesity imprints early progenitor cells potentially dictating the future metabolic status of the offspring’s adipocytes.

## INTRODUCTION

Mesenchymal stem cells (MSCs) hold a significant relevance within their cellular microenvironment [1]. Central to their significance is their paracrine activity, which orchestrates communication with the cellular milieu to maintain homeostasis [2, 3]. Referred to as the MSCs secretome, this paracrine profile consists of soluble factors such as growth factors, hormones, cytokines and metabolites, in addition to the release of extracellular vesicles [3]. These vesicles transport an array of bioactive molecules, surface receptors, and genetic information, engaging with other cells [4–8]. Thus, the secretome provides a novel approach to exploring the functional role of MSCs.

Maternal obesity presents an increased risk for the offspring of developing childhood obesity [9, 10]. However, the exact mechanisms underlying the programming of fetal adipose tissue remain elusive. Recent findings indicate that neonatal MSCs, the embryonic precursor cells for adipocytes, exhibit early signatures of an adipogenic phenotype when influenced by maternal obesity [11–14]. MSCs, as progenitor cells established during early embryonic development, are notably present in fetal tissues like the umbilical cord [15–17]. MSCs from the umbilical cord serve as an efficient model for studying the underlying mechanisms driving adipogenesis of MSCs in obese pregnancies.

Adipocyte differentiation from MSCs is accompanied by significant metabolic reprogramming, including increased mitochondrial activation and mass [18–20]. This mitochondrial activation triggers an enhanced production of reactive oxygen species (ROS), which are highly reactive oxygen-derived small molecules [21–23]. While ROS are crucial in promoting essential cell functions, including the commitment to the adipocyte lineage [19, 21, 23], levels are finely regulated by cellular detoxification systems such as antioxidant enzymes: superoxide dismutases (SOD), catalase (CAT), peroxiredoxins (PRDX) and glutathione peroxidases (GPX) [22, 23]. Furthermore, growing evidence indicates that MSCs exert antioxidant properties that function buffering ROS within their local environment [24], and facilitate intercellular communication regarding redox balance [24]. This redox secretome of MSCs aims to mitigate oxidative stress [6, 7].

In addition, MSCs undergo notable transformations during adipogenesis: they lose their fibroblastic morphology, accumulate triglycerides, and acquire mature adipocyte’s appearance and metabolic features [25]. Triglyceride accumulation is closely associated with a coordinated rise in the expression of the enzymes involved in fatty acid biosynthesis, leading to an elevated rate of de novo lipogenesis [26]. With this, lipid kinetics are a defining feature of adipocytes, and there is tight inter-regulation between adipogenic transcription and lipogenesis [26]. The secretome of MSCs includes metabolic components essential for intercellular communication regarding lipid metabolism [8]. Moreover, the secretome of adipocytes is characterized by endocrine components with metabolic functions, known as adipokines [27–29].

MSCs derived from neonates of mothers with obesity are known to have oxidative stress and higher adipogenic commitment, compared to the ones from neonates of mothers with normal weight, thereby suggesting an early programming of fetal adipose tissue [11–14, 30, 31]. We hypothesize that the impact of an obesogenic intrauterine environment on fetal MSCs might subsequently shape their paracrine redox and adipogenic activity within their surrounding environment. In this context, the present study aimed to unveil extracellular proteomic signatures that account for the early adipogenic programming of MSCs in the offspring of women with obesity.

## MATERIALS AND METHODS

### Experimental design

MSCs derived from neonates born to normal-weight mothers (NW-MSCs) and from mothers with obesity (OB-MSCs) were subjected to *in vitro* adipogenesis. The secretome of these cells was assessed at various time points: on day 0 (MSCs, progenitor cells), day 5 (early adipogenesis) and day 21 (upon reaching maturity as MSC-derived *in vitro* adipocytes, Supplementary Figure S1). The analysis focused on characterizing the secretome during the three timepoints, after which we specifically focused on proteins associated with redox mechanisms and adipogenic profile at each time point.

### Subjects

Umbilical cords were obtained from the placentas of women with obesity or normal weight, who delivered at the University Medical Centrum Groningen (UMCG) maternity ward, Groningen, The Netherlands, from July 2021 to February 2023. Umbilical cords were considered as biological waste material following routine childbirth procedures and no donor information collected. The study complied with all relevant institutional and national ethical regulations regarding the use of biological materials. A pregestational maternal body mass index (BMI) >30 kg/m² was considered for the group of women with obesity (OB, n=3) and 18.5-24.5 kg/m² for women with normal weight (NW, n=3). The inclusion criteria included women >18 years, single and term pregnancies (>37 weeks). The exclusion criteria included women with gestational diabetes, preeclampsia, preterm birth, and neonatal complications [14]. All procedures were conducted according to the Helsinki Declaration.

### Isolation of Wharton’s jelly-derived MSCs

Umbilical cords were obtained and immediately processed to get MSCs primary cultures. MSCs were isolated using the explant method [14–17]. Therefore, the umbilical cord was washed in cold Dulbecco’s Phosphate Buffered Saline (DPBS, Gibco, Thermo Fischer Scientific) and cut into three cm pieces inside a laminar flow cabinet. Each piece was cut longitudinally, and blood vessels were discarded. Wharton’s jelly explants were plated and cultured with Dulbecco’s modified Eagle medium containing 10 mM glucose (DMEM, Gibco, Thermo Fischer Scientific), 10% Fetal calf serum (FCS, Sigma-Aldrich), 5.000 UI/mL Penicillin-Streptomycin (Gibco, Thermo Fischer Scientific), and maintained at 37°C in 5% CO[. Media was changed every four days, and culture was maintained for 10-14 days. By this time, a solid population of cells had sprouted out from the explants and had covered the explant perimeter. Subsequently, the sprouted cells were trypsinized (Gibco, Thermo Fischer Scientific) and expanded into further passages by seeding at 4.000/cm² in culture flasks. The medium was changed every three days, and cells were maintained in the same culturing conditions. Cells were passaged after six days of culture (passages 1-2). Unless otherwise stated, all experiments were performed in fresh cells on passage 2.

### Cell culture and adipogenic induction

Cells were seeded at 7000 cells/cm² in 6-well plates and cultured with DMEM (Gibco, Thermo Fischer Scientific) (10 mM glucose), 10% FCS (Sigma-Aldrich), 5000 UI/ml Penicillin-Streptomycin (Gibco, Thermo Fischer Scientific) and maintained at 37°C in 5% CO2. When cells reached 80% confluency, adipogenesis was induced for 5 or 21 days with high glucose DMEM + 10% FCS, 5000 UI/ml Penicillin-Streptomycin (Gibco, Thermo Fischer Scientific), insulin (1 nM), dexamethasone (0.1 μM) and 3-isobutyl-1-methylxantine (0.5 mM) [32].

### Supernatant collection for secretome measurements and SP3 digestion

Cells were cultured in the conditions described. Upon reaching confluence, the cells were subjected to FCS starvation for 24 hours. Subsequently, the supernatant was collected and centrifuged at 400 x g for 10 minutes and stored at -80°C. Protein concentrations in all samples were determined using the bicinchoninic acid (BCA) protein assay kit (Pierce, Thermo Fisher Scientific). The further processing of the supernatants followed a SP3 digestion method [33]. Briefly, 25 μl of supernatant underwent reduction by adding dithiothreitol (10 mM) and incubated for 30 min at 57 °C. Subsequent alkylation of the sample was performed by adding iodoacetamide (30 mM) and incubating for 30 minutes at room temperature in the dark. Next, proteins were bound to prewashed Cytiva SeraMag beads (1:1 of hydrophilic and hydrophobic beads, Fisher Scientific), and diluted to 50% acetonitrile. After the removal of acetonitrile, the beads were washed with 80% (v/v) ethanol and acetonitrile. The beads were digested overnight with 40 μL of trypsin (2.5 ng/μl, Promega) at 37°C. Lastly, 10 μL of 1% (v/v) formic acid were added to stop the digestion, and untargeted proteomics analyses of the sample were further processed on the Evosep (Evosep Biosystems) [33].

### Liquid Chromatography-Mass Spectrometry Analysis

Mass spectrometry analyses were performed with a quadrupole orbitrap mass spectrometer (Orbitrap Exploris 480, Thermo Scientific). Peptide separation was achieved through liquid chromatography (Evosep One, Evosep), with a nano-LC column (EV1137 Performance column, Evosep). 10% of the sucrose fraction digests were subjected to separation utilizing the 30SPD workflow (Evosep). The mass spectrometer was configured in positive ion mode and operated in data-independent acquisition (DIA) mode. Isolation windows of 16 m/z were employed with a precursor mass range of 400-1000. Additionally, FAIMS alternated between CV -45V and -60V, incorporating three scheduled MS1 scans during each precursor mass range screening [33].

### Data analyses

Raw liquid chromatography-mass spectrometry (LC-MS) data were processed with Spectronaut (18.3.230830) (Biognosys), with the standard settings of the direct DIA workflow. MS1-level quantification was conducted using the human SwissProt database (www.uniprot.org, 20422 entries). We employed local normalization for quantification of the data, and Q-value filtering was performed using the traditional setting without imputation. Analysis of enrichment pathways was performed by Metascape considering a p-value cutoff of 0.01 and an enrichment factor of 1.5 [34]. For general characterization of the secretomes, we considered only common proteins between NW-MSCs and OB-MSCs. Further, data was transformed to log2 and Mann-Whitney U tests were applied for each protein found in the secretome of NW-MSCs and OB-MSCs, to obtain differentially regulated proteins (upregulated or downregulated proteins, cut-off log2 ratio=0.5). For downstream analysis, we considered only the differentially regulated proteins (candidate proteins).

### Enrichment pathway analyses

Candidate proteins were assessed for enrichment pathway analyses with Metascape [34], considering parental Gene Ontology pathways for comparisons of OB-MSCs versus NW-MSCs on day 0, 5 and 21 (p-value cutoff = 0.01). Analyses included Gene Ontology, Reactome, and KEGG databases.

### Redox proteins

The list of candidate proteins from days 0, 5 and 21 was evaluated in STRING databases [35] considering GO Biological Process: response to oxidative stress (GO:0006979; FDR 0.00091), regulation of cellular response to oxidative stress (GO:19000407; FDR 0.0075), response to reactive oxygen species (GO:000302; FDR 0.0096), cell redox homeostasis (GO:0045454, FDR 0.0168), glutathione metabolic process (GO:0006749; FDR 4.55e-06) and NAD metabolism (GO:0019674; FDR 0.0343).

### Adipogenic proteins

The list of candidate proteins for days 0, 5 and 21 were evaluated in STRING databases for GO Biological Process: Cellular lipid metabolic process (GO:0044255, FDR 0.0238), Triglyceride homeostasis (GO:0070328, FDR 0.0025), Lipid storage (GO:0019915; FDR 0.0049), Lipid homeostasis (GO:0.0016, FDR 0055088). Reactome/KEGG pathways: Transcriptional regulation of white adipocyte differentiation (HAS-381340; FDR 0.0329), PPAR signalling pathway (hsa033320; FDR 0.0060), Regulation of Insulin-like Growth Factor (IGF) transport proteins (HSA-381426; FDR 1.36e-20), WikiPathways: Adipogenesis (WP236; FDR 0.00265). For adipokines, proteins were identified according to reported adipose-derived secreted factors with endocrine function [27–29].

### Statistical analysis

To assess multiple comparisons, volcano plots were made using non-parametric Mann-Whitney U tests (assuming a non-parametrical distribution) with a permutation-based FDR of 0.05. Volcano plots were generated using the ggplot2 package, and heat maps were generated using the gplots package (R Studio version 3.5.1). Data was transformed to log2 for heatmaps and converted to z-score, with hierarchical clustering of proteins and samples. For antioxidant enzymes, lipid localization, fatty acid synthesis proteins and adipokines, data was expressed as median value and 25th and 75th percentiles in Graphpad Prism (GraphPad Inc.). Data was transformed to log2 and a Two-way ANOVA followed by Tukey’s range post-testing was used to compare independent variables BMI (NW-MSCs versus OB-MSCs), and days of differentiation (0, 5, 21). P values < 0.05 were considered statistically significant.

## RESULTS

### General characterization of the secretome of NW-MSCs and OB-MSCs

We analysed the proteins of the secretome of NW-MSCs and OB-MSCs. We found a total of 981 proteins on day 0, from which 96 were upregulated in OB-MSCs versus NW-MSCs (red, Figure 1A), 359 proteins were downregulated in OB-MSCs versus NW-MSCs (blue, Figure 1A), and 534 proteins that did not differ between groups (grey, Figure 1A). From the upregulated and downregulated proteins, GO analysis returned the top 3 terms: cellular process (GO:0009987), metabolic process (GO:0008152) and immune system process (GO:0002376) (red and blue, Figure 1A). For subclass analyses, the top 20 terms are detailed in Supplementary Table S1.

**Figure 1.**
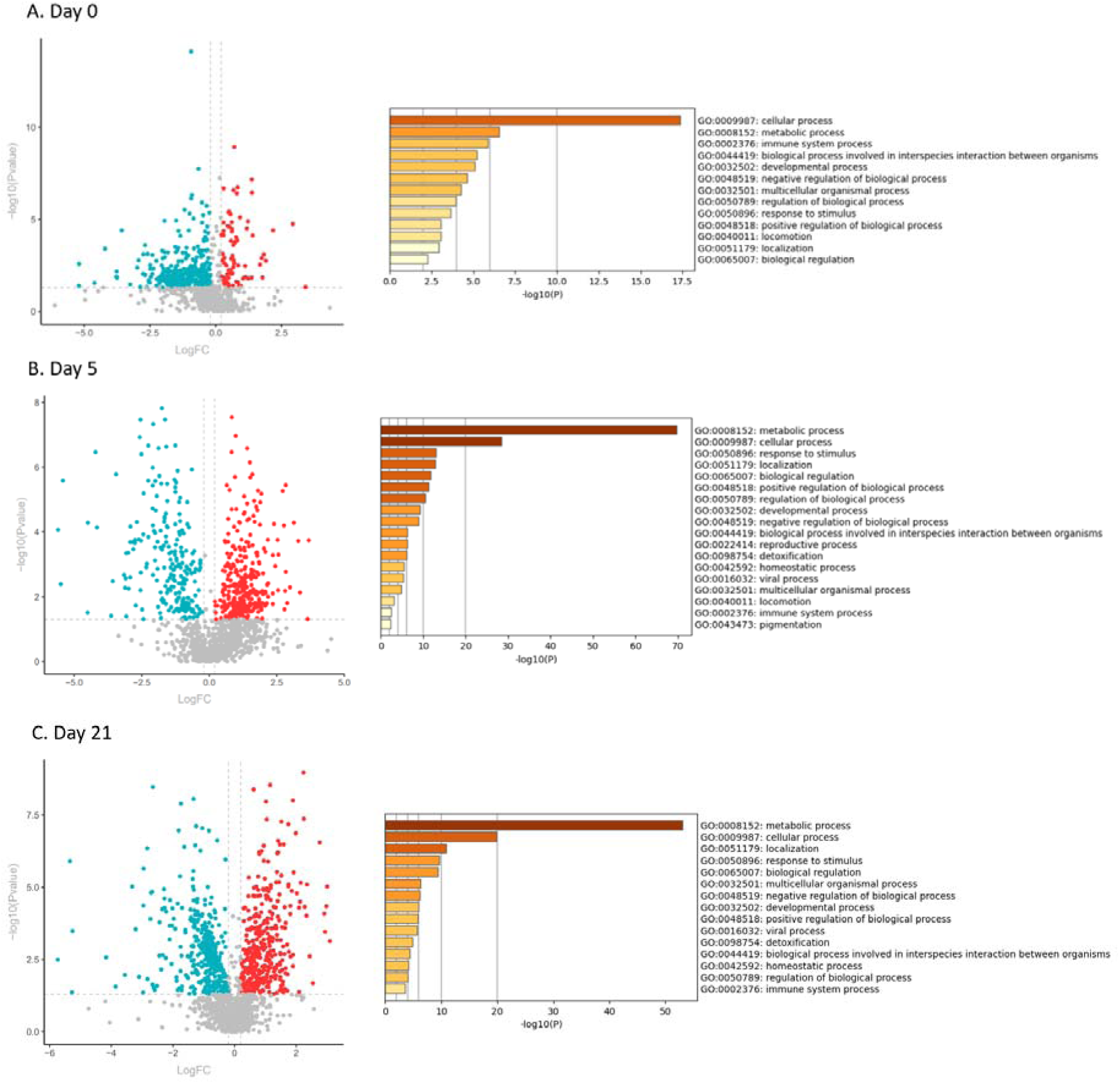
Characterization of the secretome from OB-MSCs versus NW-MSCs on days 0, 5 and 21 of adipogenesis. **A-C.** Volcano plots showing downregulation (blue), upregulation (red) and non-changed (grey) common proteins in the secretome of OB-MSCs versus NW-MSCs. Top enrichment pathways of the upregulated and downregulated proteins in the secretome of OB-MSCs versus NW-MSCs with Gene Ontology analysis.

We found 1215 proteins on day 5 in the secretome of NW-MSCs and OB-MSCs. From these proteins, 408 were upregulated (red, Figure 1B), 251 were downregulated (blue, Figure 1B), and 556 proteins did not change (grey, Figure 1B). When we evaluated the pathways represented by these upregulated and downregulated proteins, the top 3 GO terms found were: metabolic process (GO:0008152), cellular process (GO:0009987) and response to stimulus (GO: 0050896) (red and blue, Figure 1B). For subclass analyses, the top 20 terms are detailed in Supplementary Table S2.

Finally, our analyses evidenced 1506 proteins on day 21 of adipogenesis in the secretome of NW-MSCs and OB-MSCs. 331 proteins were upregulated (red, Figure 1C), 326 proteins were downregulated (blue, Figure 1C), and 849 proteins did not change (grey, Figure 1C). The top 3 GO terms associated with these proteins were: metabolic process (GO:0008152), cellular process (GO:0009987) and localization (GO: 0051179) (red and blue, Figure 1C). The top 20 terms found for subclass analyses are detailed in Supplementary Table S3.

### Redox pathways in the secretome of NW-MSCs and OB-MSCs on day 0, 5 and 21

In this study we were interested in investigating the redox profile from the proteins in the secretome of NW-MSCs and OB-MSCs on days 0, 5 and 21 of adipogenesis. The top 5 pathways represented by these candidate proteins reported response to stimulus (GO:0050896) and detoxification (GO 0098754) were found enriched on day 0, 5 and 21, suggesting a different redox secretome between both groups (Figure 1A-C). When we analysed the proteins filtered for redox analysis, we found that response to oxidative stress (GO:0006979), was the most enriched pathway represented by these proteins on day 0, 5 and 21 (Table 1).

**Table 1.** Enrichment pathways of redox proteins in the secretome of OB-MSCs versus NW-MSCs on day 0, 5 and 21 of adipogenesis.

| GO | Category | Description | Count | % | Log10(P) | Log10(q) |
| --- | --- | --- | --- | --- | --- | --- |
| **Day 0** |  |  |  |  |  |  |
| GO:0006979 | GO<br>Biological Processes | response to oxidative stress | 15 | 62.50 | -22.82 | -18.57 |
| R-HSA-9711123 | Reactome Gene Sets | Cellular response to chemical stress | 11 | 45.83 | -17.39 | -13.84 |
| GO:0042743 | GO<br>Biological Processes | hydrogen peroxide metabolic process | 7 | 29.17 | -14.78 | -11.54 |
| GO:0006749 | GO<br>Biological Processes | glutathione metabolic process | 5 | 20.83 | -9.13 | -6.17 |
| GO:1902175 | GO<br>Biological Processes | regulation of oxidative stress-induced intrinsic apoptotic signalling pathway | 4 | 16.67 | -7.28 | -4.43 |
| **Day 5** |  |  |  |  |  |  |
| GO:0006979 | GO<br>Biological Processes | response to oxidative stress | 21 | 75.00 | -34.48 | -30.23 |
| GO:1990748 | GO<br>Biological Processes | cellular detoxification | 13 | 46.43 | -24.43 | -20.48 |
| GO:0006749 | GO<br>Biological Processes | glutathione metabolic process | 11 | 39.29 | -23.20 | -19.59 |
| GO:0034599 | GO<br>Biological Processes | cellular response to oxidative stress | 11 | 39.29 | -16.31 | -13.01 |
| R-HSA-9711123 | Reactome Gene Sets | Cellular response to chemical stress | 9 | 32.14 | -12.62 | -9.54 |
| **Day 21** |  |  |  |  |  |  |
| GO:0006979 | GO<br>Biological Processes | response to oxidative stress | 12 | 85.71 | -21.14 | -16.89 |
| GO:0034599 | GO<br>Biological Processes | cellular response to oxidative stress | 8 | 57.14 | -13.67 | -9.72 |
| GO:0098869 | GO<br>Biological Processes | cellular oxidant detoxification | 5 | 35.71 | -9.37 | -5.89 |
| GO:1902175 | GO<br>Biological Processes | regulation of oxidative stress-induced intrinsic apoptotic signalling pathway | 3 | 21.43 | -5.90 | -2.90 |
| GO:0043467 | GO<br>Biological Processes | regulation of generation of precursor metabolites and energy | 3 | 21.43 | -4.47 | -1.63 |

#### 1. Redox proteins in the secretome of NW-MSCs and OB-MSCs on day 0

We evaluated the proteins particularly associated to redox mechanisms (Supplementary Table S4). On day 0, we did not identify any redox proteins increased in the secretome of OB-MSCs compared to NW-MSCs. Nevertheless, we found 23 redox proteins that showed decreased levels in the secretome of OB-MSCs compared to NW-MSCs. These decreased proteins include antioxidant enzymes such as PRDX1/3/6, CAT, as well as components of the glutathione system such as glutathione reductase (GSR) and glutathione S-transferase alpha 1 (GSTA1) (Figure 2A, Supplementary Table S4).

**Figure 2.**
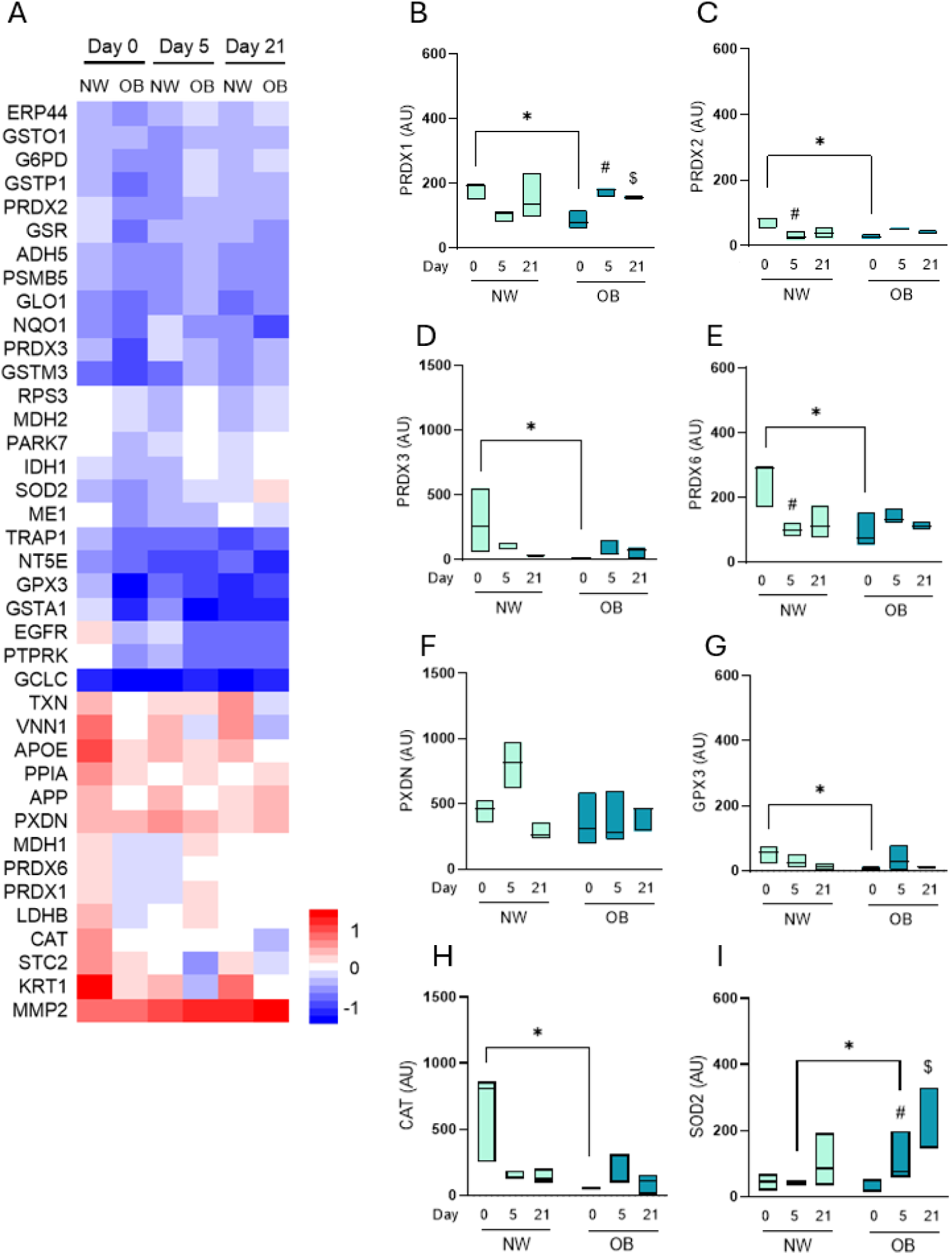
Redox proteins found in the secretome of NW-MSCs and OB-MSCs during adipogenesis. **A.** Abundance of redox proteins in the secretome of NW-MSCs and OB-MSCs for day 0, day 5 and day 21 of adipogenesis. **B-I.** Secreted antioxidant enzymes during day 0, day 5 and day 21 of adipogenesis in NW-MSCs and OB-MSCs. PRDX 1/2/3/6, peroxiredoxins; PXDN, peroxidasin; GPX3, glutathione peroxidase 3; CAT, catalase; SOD2, superoxide dismutase 2. Two-way ANOVA: Effect of day of differentiation over the expression of SOD2 (p=0.006). Effect of obesity over the expression of PRDX3 and CAT (p=0.04 and 0.007, respectively). Tukey’s range post testing: *p<0.05 (NW-MSCs versus OB-MSCs); #p<0.05 (day 5 versus day 0). $p<0.05 (day 21 versus day 0).

#### 2. Redox proteins in the secretome of NW-MSCs and OB-MSCs on day 5 (early adipogenesis)

On day 5 of adipogenesis, we identified 18 redox proteins that were increased in the secretome of OB-MSCs compared to NW-MSCs. These proteins included antioxidant enzymes such as: superoxide dismutase 2 (SOD2), PRDX1/2/6, and components of the glutathione system such as glutathione S-transferase P1 (GSTP1) and glutathione S-transferase O1 (GSTO1), among others redox mediators. Conversely, only four proteins were shown to be decreased in the secretome of OB-MSCs compared to NW-MSCs: peroxidasin homolog (PXDN), apolipoprotein E (APOE), stanniocalcin 2 (STC2) and keratin 1 (KRT1) (Figure 2A, Supplementary Table S4).

#### 3. Redox proteins in the secretome of NW-MSCs and OB-MSCs on day 21 (adipocytes)

On day 21 of adipogenesis, we identified seven redox proteins that were more abundant in the secretome of OB-MSCs compared to NW-MSCs, including PRDX6 and PXDN. We found four redox proteins decreased in the secretome of OB-MSCs compared to NW-MSCs: protein receptor-type tyrosine-protein phosphatase kappa (PTPRK), pantetheinase (VNN1), NAD(P)H dehydrogenase quinone 1 (NQO1) and KRT1 (Figure 2A, Supplementary Table S4).

### Antioxidant enzymes in the secretome of NW-MSCs and OB-MSCs during adipogenesis

As indicated above, we found many antioxidant enzymes to be changed in the secretome of NW-MSCs versus OB-MSCs (Figure 2A). We therefore analysed the presence of these enzymes throughout adipogenesis (Figure 2B-I). There was an effect of day of differentiation over the expression levels of SOD2 (p=0.006). Further, antioxidant enzymes PRDX2 and PRDX6 were decreased on day 5 compared to day 0 for NW-MSCs (p<0.05, Figure 2C and 2E). On the contrary, for OB-MSCs, PRDX1 and SOD2 were increased on days 5 and 21 compared to day 0 (p<0.05, Figure 2B and 2I).

Next, our analyses showed an effect of obesity on expression levels of PRDX3 and CAT (p=0.04 and 0.007, respectively). Further, on day 0, many of the antioxidant enzymes were less abundant in the secretome of OB-MSCs compared to NW-MSCs: PRDX1/2/3/6, GPX3 and CAT (p<0.05, Figure 2B-E, and 2G-H). Interestingly, SOD2 secretion was increased the secretome of OB-MSCs compared to NW-MSCs on day 5 (p<0.05, Figure 2I).

### Adipogenic pathways in the secretome of NW-MSCs and OB-MSCs on days 0, 5 and 21

We were interested in investigating the adipogenic proteins in the secretome of NW-MSCs and OB-MSCs on day 0, 5 and 21 of adipogenesis. We found that cellular lipid catabolic process (GO:0044242) and lipid localization (GO:0010876) were the most enriched pathways represented on days 0, 5 and 21 (Table 2).

**Table 2.**
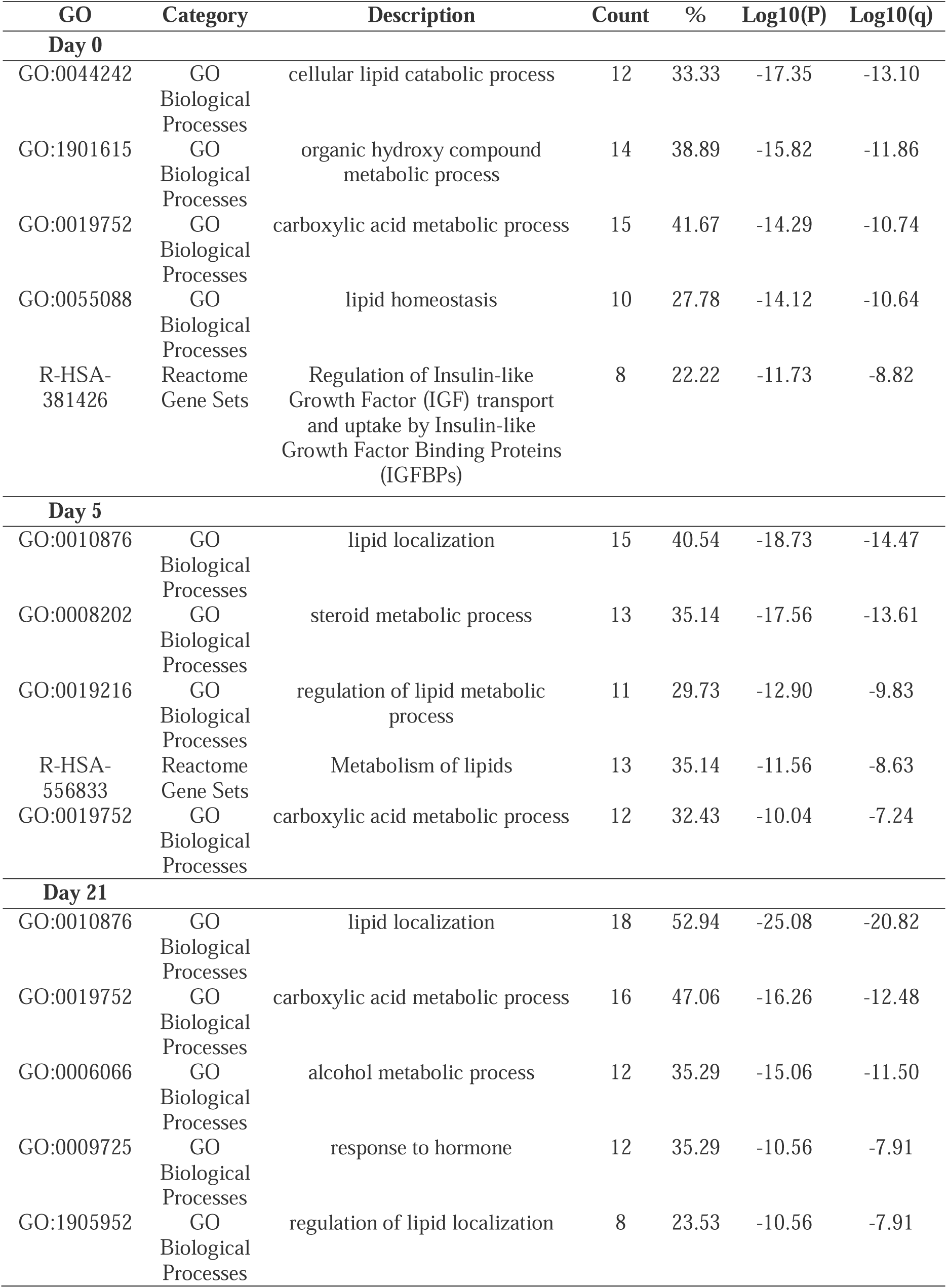
Enrichment pathways of adipogenic proteins in the secretome of OB-MSCs versus NW-MSCs on day 0, 5 and 21 of adipogenesis.

| GO | Category | Description | Count | % | Log10(P) | Log10(q) |
| --- | --- | --- | --- | --- | --- | --- |
| **Day 0** |  |  |  |  |  |  |
| GO:0044242 | GO<br>Biological Processes | cellular lipid catabolic process | 12 | 33.33 | -17.35 | -13.10 |
| GO:1901615 | GO<br>Biological Processes | organic hydroxy compound metabolic process | 14 | 38.89 | -15.82 | -11.86 |
| GO:0019752 | GO<br>Biological Processes | carboxylic acid metabolic process | 15 | 41.67 | -14.29 | -10.74 |
| GO:0055088 | GO<br>Biological Processes | lipid homeostasis | 10 | 27.78 | -14.12 | -10.64 |
| R-HSA-381426 | Reactome Gene Sets | Regulation of Insulin-like Growth Factor (IGF) transport and uptake by Insulin-like Growth Factor Binding Proteins (IGFBPs) | 8 | 22.22 | -11.73 | -8.82 |
| **Day 5** |  |  |  |  |  |  |
| GO:0010876 | GO<br>Biological Processes | lipid localization | 15 | 40.54 | -18.73 | -14.47 |
| GO:0008202 | GO<br>Biological Processes | steroid metabolic process | 13 | 35.14 | -17.56 | -13.61 |
| GO:0019216 | GO<br>Biological Processes | regulation of lipid metabolic process | 11 | 29.73 | -12.90 | -9.83 |
| R-HSA-556833 | Reactome Gene Sets | Metabolism of lipids | 13 | 35.14 | -11.56 | -8.63 |
| GO:0019752 | GO<br>Biological Processes | carboxylic acid metabolic process | 12 | 32.43 | -10.04 | -7.24 |
| **Day 21** |  |  |  |  |  |  |
| GO:0010876 | GO<br>Biological Processes | lipid localization | 18 | 52.94 | -25.08 | -20.82 |
| GO:0019752 | GO<br>Biological Processes | carboxylic acid metabolic process | 16 | 47.06 | -16.26 | -12.48 |
| GO:0006066 | GO<br>Biological Processes | alcohol metabolic process | 12 | 35.29 | -15.06 | -11.50 |
| GO:0009725 | GO<br>Biological Processes | response to hormone | 12 | 35.29 | -10.56 | -7.91 |
| GO:1905952 | GO<br>Biological Processes | regulation of lipid localization | 8 | 23.53 | -10.56 | -7.91 |

#### 1. Adipogenic proteins in the secretome of NW-MSCs and OB-MSCs on day 0

In our analysis we identified four adipogenic proteins that were increased in the secretome of OB-MSCs compared to NW-MSCs on day 0: lysosomal acid glucosylceramidase (GBA1), macrophage migration inhibitory factor (MIF), trifunctional enzyme subunit beta (HADHB) and insulin-like growth factor binding protein 6 (IGFBP6). Conversely, 32 proteins were decreased in the secretome of OB-MSCs compared to NW-MSCs, including proteins involved in lipid metabolism (APOE; aldo-keto reductase 1 A1, AKR1A1; and prostaglandin reductase 1, PTGR1) and regulators of insulin pathway (insulin-like growth factor binding protein acid labile subunit, IGFALS; insulin-like growth factor-binding protein 2, IGFBP2), among others (Figure 3A, Supplementary Table S5).

**Figure 3.**
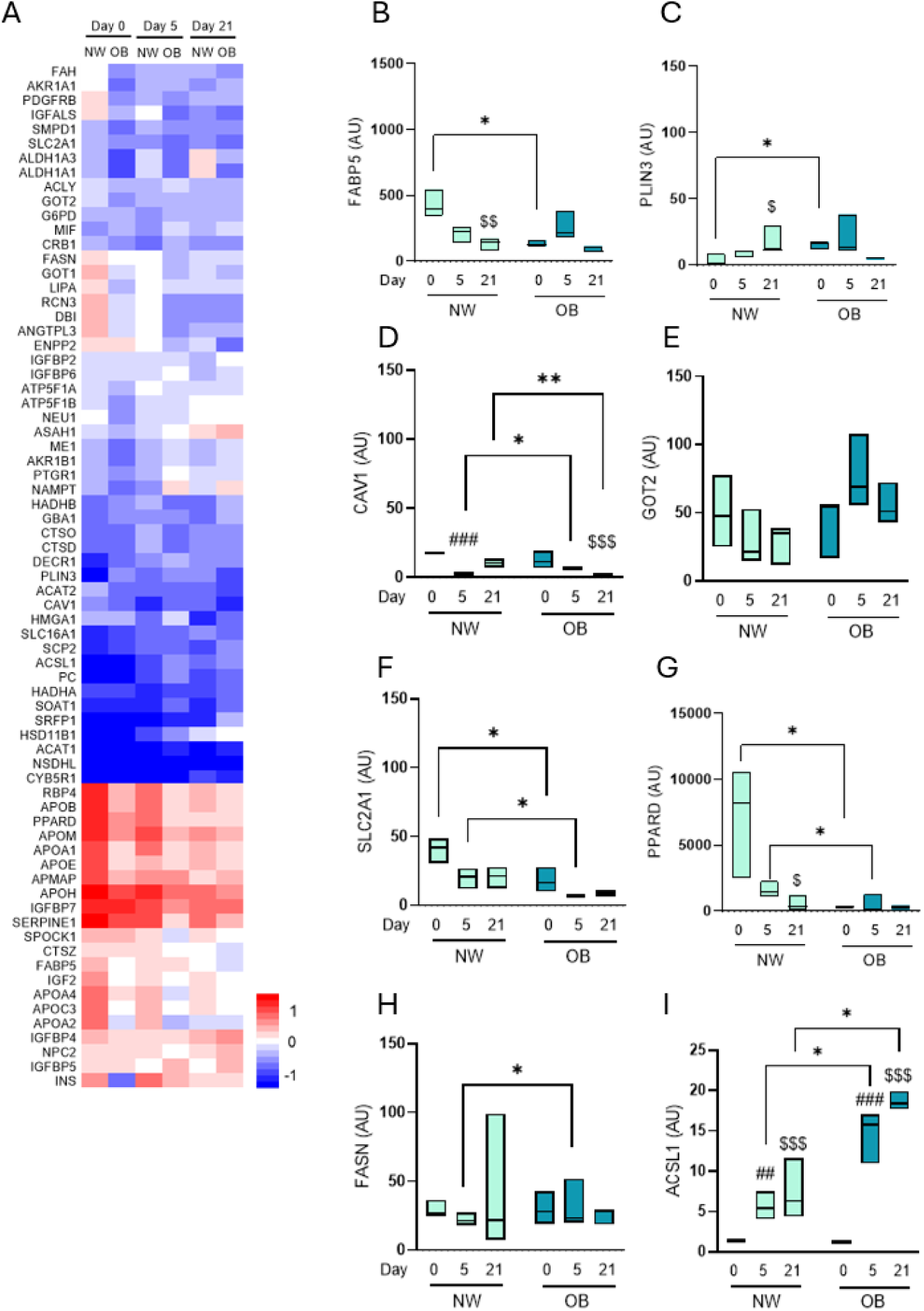
Adipogenic proteins in the secretome of NW-MSCs and OB-MSCs during adipogenesis. **A.** Abundance of adipogenic proteins in the secretome of NW-MSCs and OB-MSCs for day 0, 5 and 21 of adipogenesis. **B-G.** Lipid localization proteins. FABP5, fatty acid binding protein 5; PLIN3, perilipin 3; CAV1, caveolin 1, ANGPTL3, angiopoietin like protein 3; GOT2, Glutamic-oxaloacetic transaminase; SLC2A1, soluble carrier family 2 member 1; PPARD, peroxisome proliferator-activator receptor delta. Two-way ANOVA: Effect of day of differentiation on FABP5, CAV1 and SLC2A1 (p=0.001, 0.0001 and 0.04, respectively). Effect of obesity on FABP5, SLC2A1 and PPARD (p=0.01, 0.0004 and 0.0006, respectively). **H-I.** Fatty acid synthesis proteins. FASN, fatty acid synthase; ACSL1, acyl-COA synthase long chain 1. Two-way ANOVA: Effect of day of differentiation on ACSL1 (p=0.0001). Effect of obesity on ACSL1 (p=0.003). Tukey’s range post testing: *p<0.05 (NW-MSCs versus OB-MSCs); **p<0.01 (NW-MSCs versus OB-MSCs); ##p<0.01 (day 5 versus day 0); ###p<0.001 (day 5 versus day 0); $p<0.05 (day 21 versus day 0); $$p<0.01 (day 21 versus day 0); $$$p<0.001 (day 21 versus day 0).

#### 2. Adipogenic proteins in the secretome of NW-MSCs and OB-MSCs on day 5 (early adipogenesis)

On day 5, we identified 18 proteins with higher abundance in the secretome of OB-MSCs compared to NW-MSCs. These proteins were mainly related to lipid metabolism (visfatin /nicotinamide phosphoribosyltransferase, NAMPT; glucose-6-phosphate 1-dehydrogenase, G6PD), adipogenesis (high mobility group protein A1, HMGA1; and MIF), among other mechanisms. In contrast, 17 proteins were found with lower abundance in the secretome of OB-MSCs compared to NW-MSCs: proteins related to lipid metabolism (apolipoprotein A1, APOA2; and apolipoprotein A1, APOA1), regulation of insulin pathway (insulin, INS; and insulin-like growth factor binding protein 6 and 7, IGFBP6/7), and adipogenesis (testican 1, SPOCK1; and solute carrier 2 facilitated glucose transporter 1/GLUT1, SLC2A1) (Figure 3A, Supplementary Table S5).

#### 3. Adipogenic proteins in the secretome of NW-MSCs and OB-MSCs on day 21 (adipocytes)

On day 21, our analyses revealed 16 proteins that were increased in the secretome of OB-MSCs compared to NW-MSCs, which were mainly related to lipid metabolism (sterol O-acyltransferase 1, SOAT1; and HADHB). Conversely, 18 proteins were decreased in the secretome of OB-MSCs compared to NW-MSCs, also related to lipid metabolism (aldehyde dehydrogenase 1 A1, ADH1A1; and caveolin-1, CAV1) (Figure 3A, Supplementary Table S5).

### Proteins related to lipid localization in the secretome of NW-MSCs and OB-MSCs during adipogenesis

As indicated above, we identified lipid localization (GO: 0010876) as the most enriched pathway from the adipogenic proteins in the secretome of OB-MSCs and NW-MSCs (day 5 and 21, Table 2). Consequently, we evaluated the temporal expression of these proteins during adipogenesis. Our analysis showed an effect of day of differentiation on the expression of fatty acid binding protein 5 (FABP5), CAV1 and SLC2A1 (p=0.001, 0.0001 and 0.04, respectively). Moreover, we evidenced that FABP5 was decreased on day 21 compared to day 0 in NW-MSC secretome (p<0.01), while PLIN3 was increased on day 21 compared to day 0 for NW-MSCs (p<0.05, Figure 3B-C). This was not seen in OB-MSCs (Figure 3B-C). Further, CAV1 levels were decreased on day 5 compared to day 0 in NW-MSCs (P=0.001), while it was decreased on day 21 compared to day 0 in OB-MSCs (p<0.0001, Figure 3D). Peroxisome proliferator-activated receptor delta (PPARD) was decreased on day 21 compared to day 0 in NW-MSCs (p=0.0001, Figure 3D and 3G).

Next, we found an effect of obesity over the expression of FABP5, SLC2A1, and PPARD (p=0.01, 0.0004 and 0.0006, respectively). On day 0, FABP5 was lower (p<0.05, Figure 3B), while PLIN3 levels were higher in the secretome of OB-MSCs versus NW-MSCs (p<0.05, Figure 3C). CAV1 was higher in the secretome of OB-MSCs compared to NW-MSCs on day 5 but decreased on day 21 in OB-MSCs compared to NW-MSCs secretome (p<0.05 and <0.001, respectively, Figure 3D). SLC2A1 and PPARD levels were decreased in OB-MSCs compared to NW-MSCs on day 0 and day 5 (p<0.05, Figure 3F and 3G).

### Fatty acid synthesis in the secretome of NW-MSCs and OB-MSCs during adipogenesis

We focused on the secreted proteins related to fatty acid synthesis that could influence lipid accumulation during adipogenesis. Our analyses showed an effect of day of differentiation (p=0.0001) over levels of Long-chain-fatty-acid--CoA ligase 1 (ACSL1), but no effect over fatty acid synthase (FASN) levels (p=0.08). Further, we found that ACSL1 was increased during adipogenesis (day 5 and 21 compared to day 0) for both NW-MSCs and OB-MSCs (p<0.01 and p<0.001).

Next, our analyses showed an effect of obesity (0,003) over levels of ACSL1. Further, our results showed that fatty acid synthase (FASN) levels were higher in the secretome of OB-MSCs compared to NW-MSCs on day 5 (p<0.05, Figure 3H). There was a higher abundance of ACSL1 in the secretome of OB-MSCs compared to NW-MSCs on day 5 and day 21 (p<0.05, Figure 3I).

### Adipokines in the secretome of NW-MSCs and OB-MSCs during adipogenesis

We identified the presence of retinol-binding protein 4 (RBP4), visfatin/NAMPT, plasminogen activator inhibitor 1 (PAI-1), and the chemokines C-X-C motif chemokine 1 (CXCL1) and complement C1q tumour necrosis factor-related protein 1 (C1QTNF1) (Figure 4A). We also found novel adipokines, including cathepsins, angiopoietin-related protein 3 (ANGPTL3), secreted frizzled-related protein 1 (SFRP1), and MIF (Figure 4A). Our analyses showed an effect of day of differentiation on expression levels of visfatin/NAMPT, PAI-1, ANGPTL3, SFRP1 and Cathepsin Z (p=0.0007, 0.009, 0.004, 0.0015 and 0.0015, respectively). Further, our results indicated higher levels of visfatin/NAMPT and C1qTNF1 on day 5 and 21 compared to day 0 for OB-MSCs (p<0.01 and <0.05, Figure 4C and 4L), while higher levels of SFRP1 on day 21 compared to day 0 (p<0.001, Figure 4F). Contrarywise, OB-MSCs showed lower levels of PAI-1 on day 5 compared to day 0 (p<0.05, Figure 4D) and Cathepsin Z (p<0.05 and <0.01, respectively day 5 and day 21 compared to day 0, Figure 4I).

**Figure 4.**
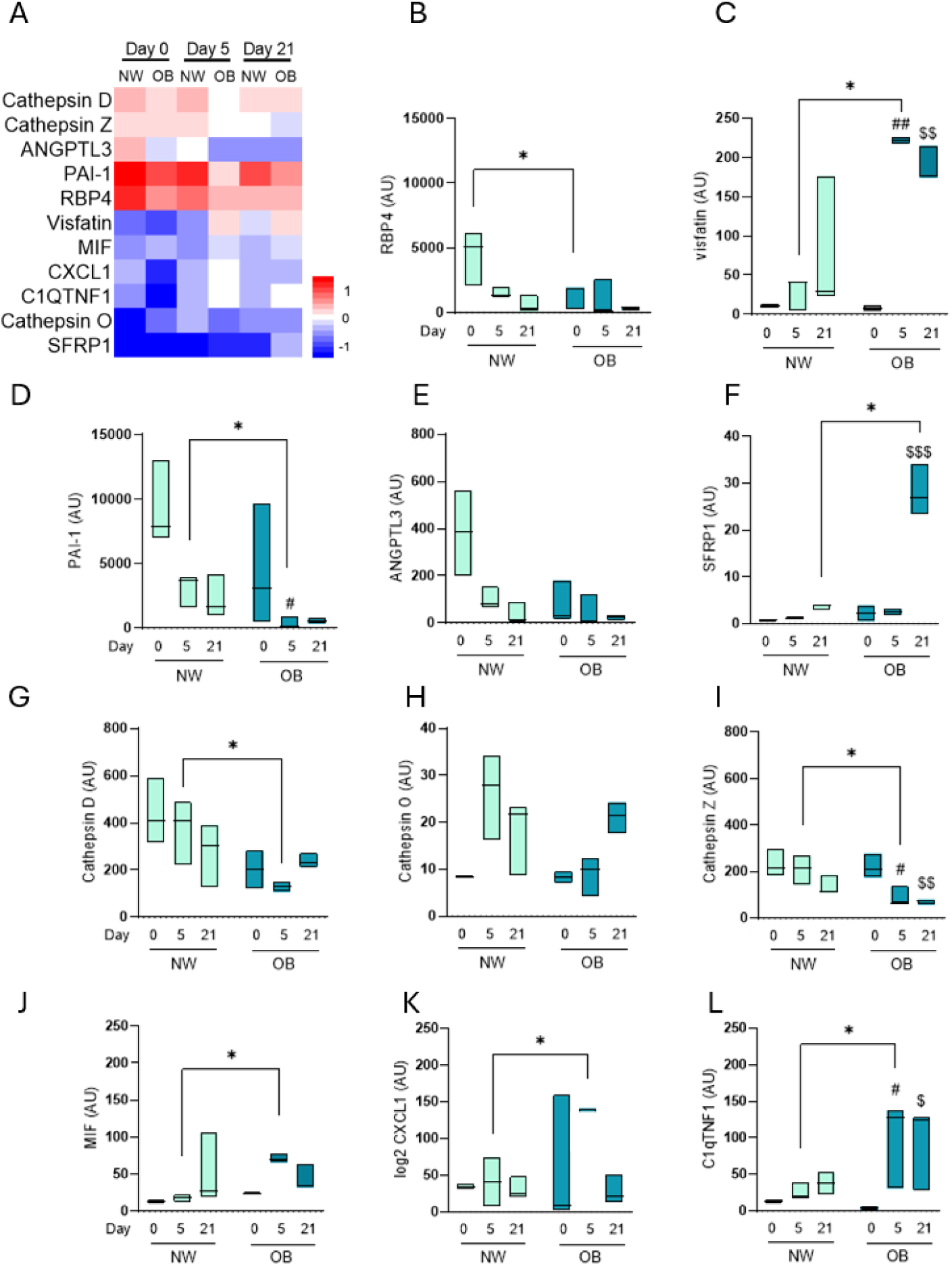
Adipokines found in the secretome of NW-MSCs and OB-MSCs during adipogenesis. **A.** Abundance of secreted adipokines in the secretome of NW-MSCs and OB-MSCs for day 0, day 5 and day 21 of adipogenesis. **B-L.** Secreted adipokines in NW-MSCs and OB-MSCs for day 0, day 5 and day 21 of adipogenesis. RBP4, retinol-binding protein 4; ANGPTL3, angiopoietin-like protein 3; SFRP1, secreted frizzle related protein 1; MIF, macrophage inhibitory factor; PAI-1, plasminogen activator inhibitor 1; CXCL1, chemokine ligand 1; C1QTNF1, C1q tumour necrosis factor related protein 1. Two-way ANOVA: Effect of day of differentiation over levels of visfatin/NAMPT, PAI-1, ANGPTL3, SFRP1 and Cathepsin Z (p=0.0007, 0.009, 0.004, 0.0015 and 0.0015, respectively). Effect of obesity over the expression of RBP4, visfatin/NAMPT, PAI-1, ANGPTL3, SFRP1, Cathepsin D, Cathepsin Z and MIF (p= 0.03, 0.01, 0.001, 0.003, 0.03, 0.004, 0.003 and 0.01, respectively). Tukey’s post testing: *p<0.05 (NW-MSCs versus OB-MSCs); #p<0.05 (day 5 versus day 0); ##p<0.01 (day 5 versus day 0); $p<0.05 (day 21 versus day 0); $$p<0.01 (day 21 versus day 0); $$$p<0.001 (day 21 versus day 0).

Further, our results showed an effect of obesity over the expression of RBP4, visfatin/NAMPT, PAI-1, ANGPTL3, SFRP1, Cathepsin D, Cathepsin Z and MIF (p=0.03, 0.01, 0.001, 0.003, 0.03, 0.004, 0.003 and 0.01, respectively). We evidenced lower RBP4 levels in OB-MSCs versus NW-MSCs secretome on day 0 (p<0.05, Figure 4B). Interestingly, on day 5, visfatin/NAMPT, MIF, CXCL1 and C1QTNF1 were increased, while PAI-1, cathepsins D and Z were decreased in the secretome of OB-MSCs compared to NW-MSCs (p<0.05, Figure 4C, 4J-L, 4D). On day 21, SFRP1 levels were increased in the secretome of OB-MSCs compared to NW-MSCs (p<0.05, Figure 4F). Notably, we found the presence of interleukin 6 (IL-6) in the secretome of OB-MSCs on day 0, while no expression was detected in NW-MSCs (data not shown).

## DISCUSSION

In this study, we characterized the secretome of NW-MSCs and OB-MSCs. Our analysis revealed fundamental differences associated with cellular and metabolic processes. Moreover, the secretome of OB-MSCs exhibited lower expression of proteins related to redox and lipid localization, while showing higher expression of proteins associated with lipid synthesis and proinflammatory adipokines, compared to NW-MSCs. The secretion of signalling factors is a pivotal function of MSCs, and proteomic analyses of the secretome offers crucial insights into alterations in cell signalling and intercellular communication [36–38].

Our results show that redox proteins are decreased in the secretome of OB-MSCs compared to NW-MSCs in the basal state (day 0). Thus, it suggests a decreased antioxidant potential of OB-MSCs within the extracellular space, compared to NW-MSCs, and could interfere to preserve the tissue’s health and redox communication with the cellular milieu [39]. Our findings align with animal studies reporting decreased levels of redox proteins in the secretome of bone marrow-derived MSCs from high-fat diet mice. In this study, they found an absence of glutamate-cysteine ligase (GCL), PRDX5 and PRDX6 compared to normal diet mice [38]. These observations collectively indicate that maternal obesity disrupts the redox homeostasis of their offspring’s MSCs within the extracellular environment [14].

We observed that OB-MSCs show an increased secretion of redox proteins during day 5 and 21 of adipogenic differentiation, compared to NW-MSCs. These results can be related to the fact that mitochondrial metabolism and oxidative stress have been implicated in the differentiation of MSCs and adipogenic commitment [19], which is associated to an activation of antioxidant mechanisms [39–40]. Among the redox proteins in the secretome between NW-MSCs and OB-MSCs, we found that Parkinson’s disease protein 7 (PARK7) and mitochondrial heat shock protein 75 kDa (TRAP1) were significantly decreased on day 0 but increased on day 5 and 21 of adipogenesis in OB-MSCs versus NW-MSCs. PARK7 is known to act as a redox sensor and protection against oxidative stress through ERK1/2 activation [41], while TRAP1 is a mitochondrial chaperone protein that has been reported to maintain mitochondrial homeostasis [42]. This suggests that PARK7 and TRAP1 play an important role in regulating mitochondrial ROS levels during adipogenesis in the context of obesity, and further studies should unveil the intracellular activity of these proteins during adipogenesis.

In this study, we found that secreted SOD2 did not change during adipogenesis for NW-MSCs. However, it was increased on days 5 and 21 as compared to day 0 in the secretome of OB-MSCs. SOD2 is a mitochondrial antioxidant enzyme known to be upregulated during adipogenesis [43, 44]. Moreover, it has been described that under oxidative stress, cells are able to excrete mitochondrial cargo material through extracellular vesicles [45, 46]. Therefore, our findings suggest that OB-MSCs are under oxidative stress during adipogenic commitment, which could explain the higher secretion of SOD2 into the extracellular space.

We found lower PRDXs (PRDX1, PRDX2, PRDX3 and PRDX6) in the secretome of OB-MSCs versus NW-MSCs on day 0, suggesting a lower antioxidant capacity. Peroxiredoxins are a family of cysteine-dependent peroxidase enzymes that regulate peroxide levels [47] and their secretion has been described as a protective mechanism within the MSCs environment [37, 38], therefore, OB-MSCs could have a less efficient antioxidant system compared to NW-MSCs. It is noteworthy to mention that PRDX2 and PRDX6 were decreased on day 5 compared to day 0 in the secretome of NW-MSCs, while this was not evidenced in OB-MSCs. This observation could be explained by the fact that MSCs are known to have important detoxification properties within their extracellular milieu [24, 45, 46], while mature adipocytes do not have this function. Therefore, this paracrine capacity could be reduced in healthy preadipocytes, and adipocytes (day 5 and day 21) compared to healthy MSCs (day 0) in NW-MSCs [48]. On the contrary, we observed a higher secretion of PRDX1 on day 5 compared to day 0 in OB-MSCs. This suggests that the lower levels of peroxiredoxins in OB-MSCs compared to NW-MSCs on day 0, could be compensated during early adipogenesis in OB-MSCs, thereby enhancing antioxidant capacity during adipogenic commitment.

Major differences in proteins related to lipid localization were found between NW-MSCs and OB-MSCs secretomes, while adipogenesis *per se* did not affect these proteins. We found that PPARD was decreased in the secretome of OB-MSCs compared to NW-MSCs. PPARD is a nuclear receptor that plays an important role in lipid metabolism by promoting fatty acid oxidation [49], and the overexpression of PPARD has been associated with a decrease in body weight, adipocyte triglyceride accumulation, circulating free fatty acids and circulating triglyceride levels [50]. Thus, our results suggest a possible increase in triglyceride accumulation in MSCs and preadipocytes, and future studies should evaluate the intracellular expression of PPARD. This is in line with the higher abundance of proteins related to fatty acid synthesis in the secretome of OB-MSCs: we found higher secretion of FASN and ACSL1 compared to NW-MSCs on days 5 and 21, indicating higher cellular lipid synthesis [51, 52]. It is also in line with higher adipogenic potential of OB-MSCs as shown by the higher lipid accumulation, PPARγ expression, bigger lipid droplet size and adipocyte hypertrophy [11–13, 31]. Collectively, these data suggest an altered lipid metabolism in OB-MSCs compared to NW-MSCs during adipogenesis and highlights potential targets in the secretome for further investigation into the mechanisms underlying metabolic dysfunction in the progeny.

In our study, we aimed to evaluate the secretion of adipokines during adipogenesis. Our analysis did not detect classic adipokines such as leptin and adiponectin, while other studies show controversial results on their presence, mainly attributed to technical approach [48, 53–56]. This discrepancy highlights the need to consider the source of MSCs and methodologies used for adipogenic differentiation, when addressing these findings. We did identify novel adipokines: SFRP1 was more abundant in the secretome of OB-MSCs compared to NW-MSCs on day 21. SFRP1 is a proadipogenic peptide that inhibits the Wnt/β-catenin signalling pathway and is upregulated in the early stages of obesity [57]. On day 5, considered an early adipogenic stage, we observed higher levels of visfatin/NAMPT, MIF, CXCL1 and C1qTNF1 in the secretome of OB-MSCs compared to NW-MSCs. These adipokines are proinflammatory and correlate with obesity and type II diabetes [58–61]. In general, the dynamic regulation of these adipokines during adipogenesis indicate a potential proinflammatory state of adipose tissue and could affect metabolic regulation [62, 63]. It is important to note that MSCs are present in neonatal adipose tissue and later in life, therefore, the proinflammatory secretome of OB-MSCs could contribute to metabolic dysfunction in postnatal life.

In conclusion, this study unveiled differences in redox and adipogenic proteins within the secretome of OB-MSCs and NW-MSCs. While maternal obesity led to a reduction in redox capacity in the secretome of OB-MSCs compared to NW-MSCs on day 0, adipogenesis triggered many redox proteins increase in the secretome of OB-MSCs compared to NW-MSCs. Noteworthy differences were also observed in the secretion of lipid synthesis proteins and proinflammatory adipokines secreted by OB-MSCs compared to NW-MSCs. These events may contribute to the dysregulated adipogenesis observed in obesity. We propose that the study of the secretome of these cells provide insights into the programming of adipose tissue, while underlying the premise that maternal obesity can influence early progenitor cells and dictate the future metabolic status of adipocytes.

## ACKNOWLEDGEMENTS

We thank the Interfaculty Mass Spectrometry Center and Dr. J.C Wolters for the valuable assistance regarding sample processing and primary analyses.

## DATA AVAILABILITY

Data will be made available on request.

## AUTHOR CONTRIBUTIONS

SB: Conceptualization, methodology, investigation, formal analysis, writing. EPM: Conceptualization, methodology, formal analysis. PC: Conceptualization. MMF: Conceptualization, review and editing. TP: Review and editing.

## Supplementary Material

**Supplementary Figure S1.**
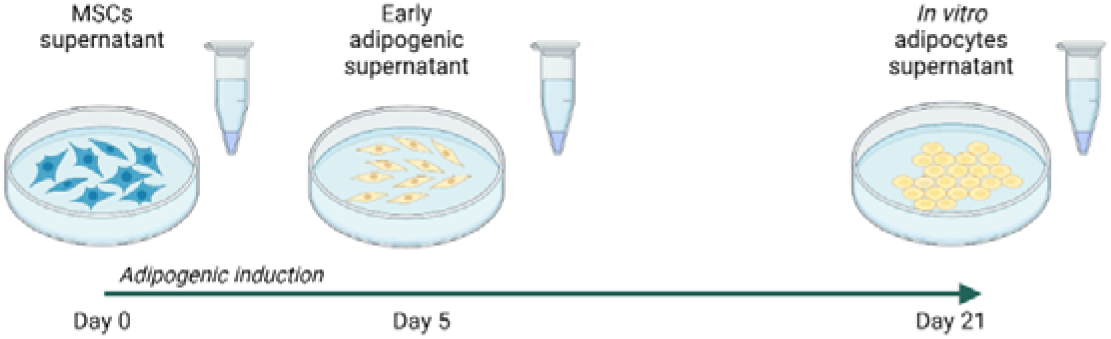
Illustrative representation of the experimental design. NW-MSCs and OB-MSCs were induced for *in vitro* adipogenesis for 21 days. Supernatant was collected on day 0, 5 and 21 for secretomic analysis by LC-MS. Created in BioRender.com

**Supplementary Table S1.**
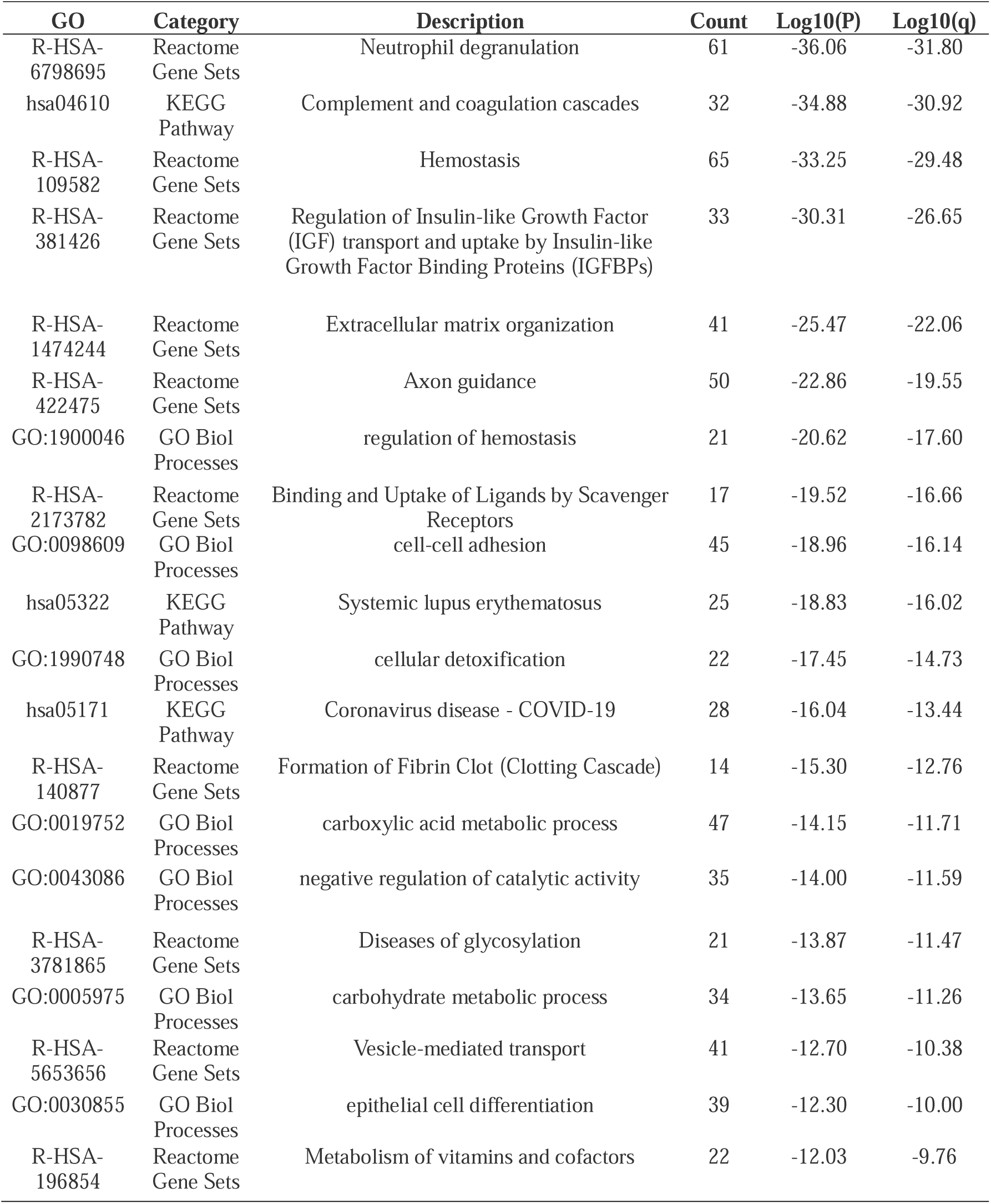
Top 20 subclass enrichment pathways of OB-MSCs versus NW-MSCs on day 0.

**Supplementary Table S2.**
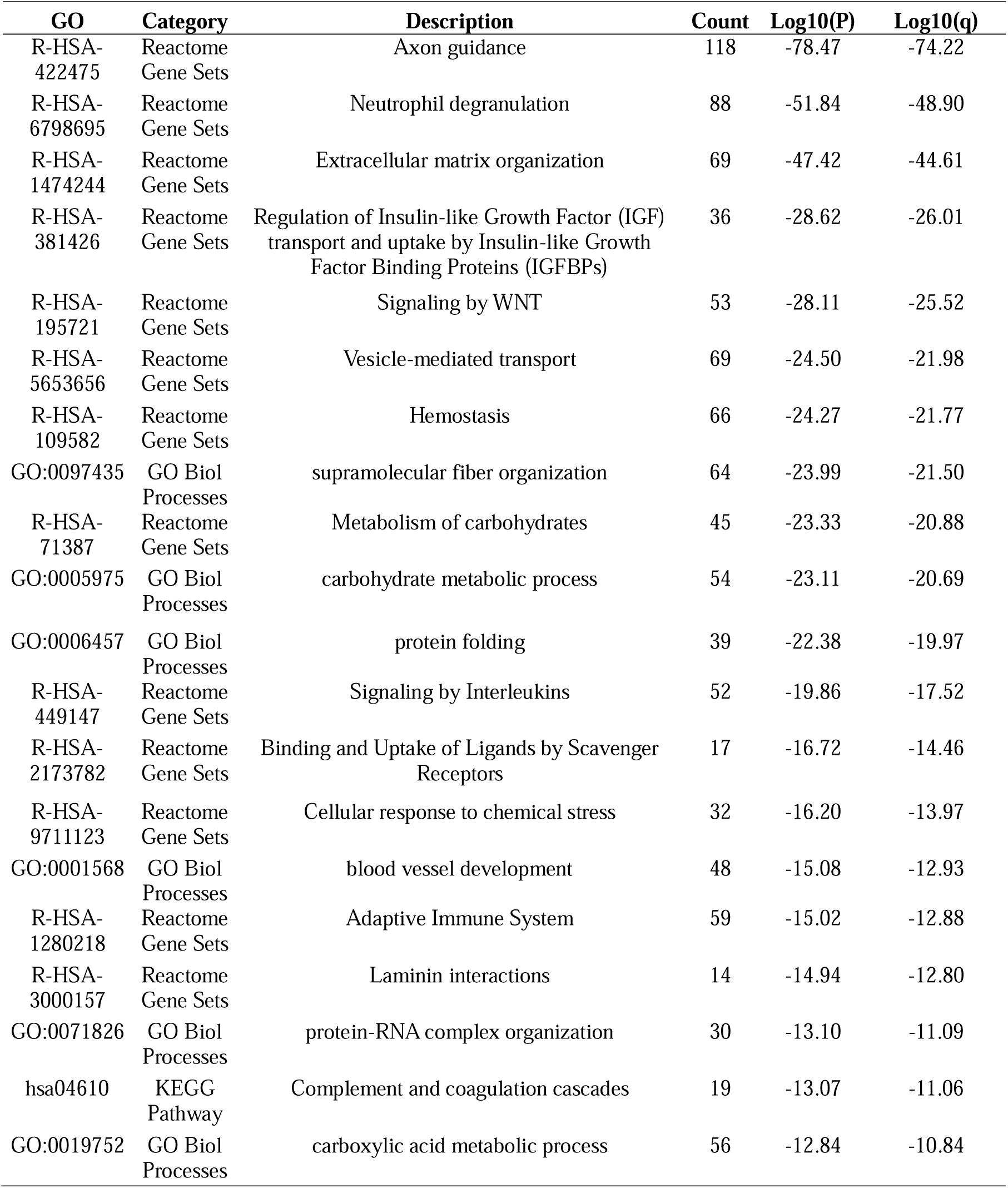
Top 20 subclass enrichment pathways of OB-MSCs versus NW-MSCs on day 5.

**Supplementary Table S3.**
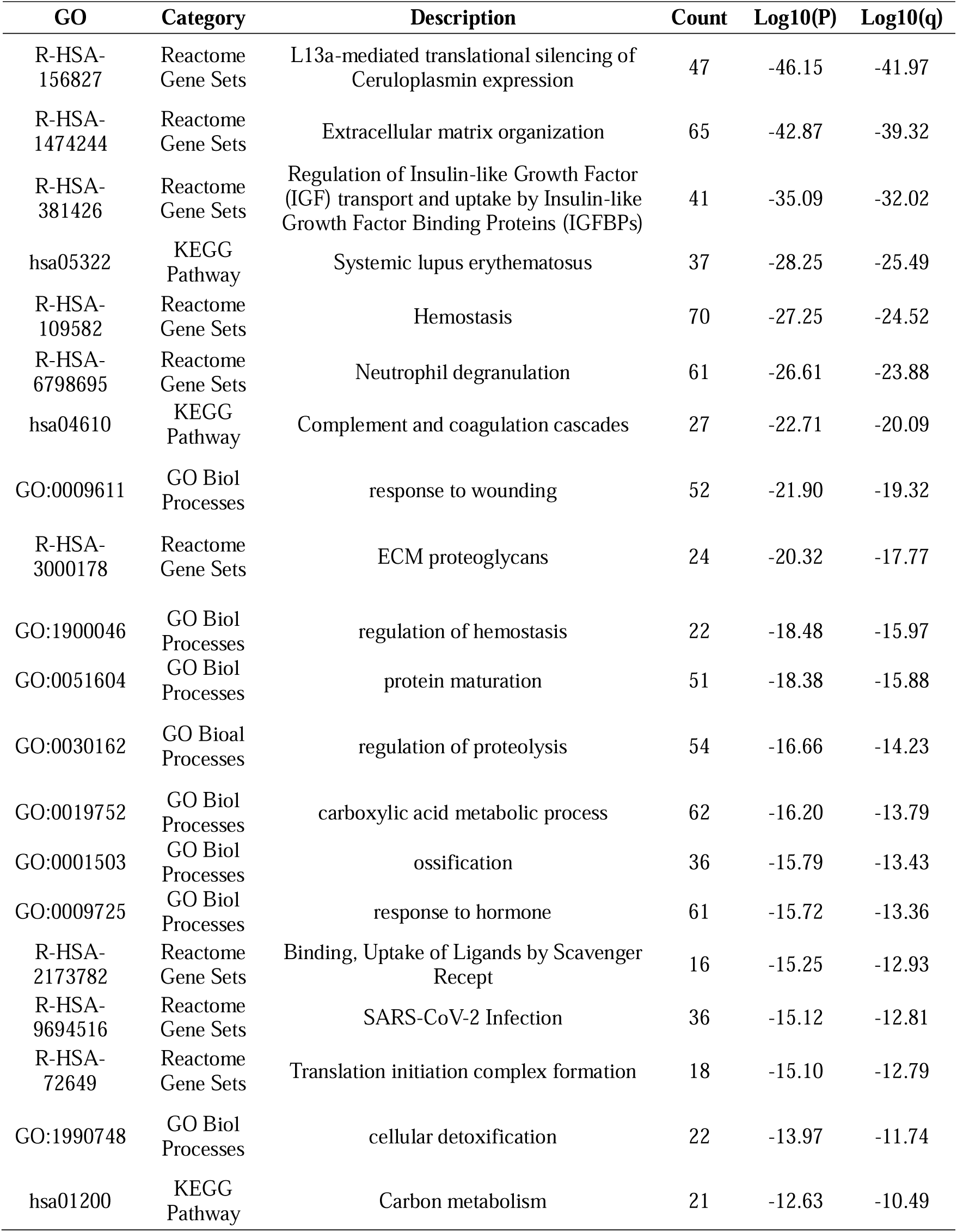
Top 20 subclass enrichment pathways of OB-MSCs versus NW-MSCs on day 21.

**Supplementary Table S4.**
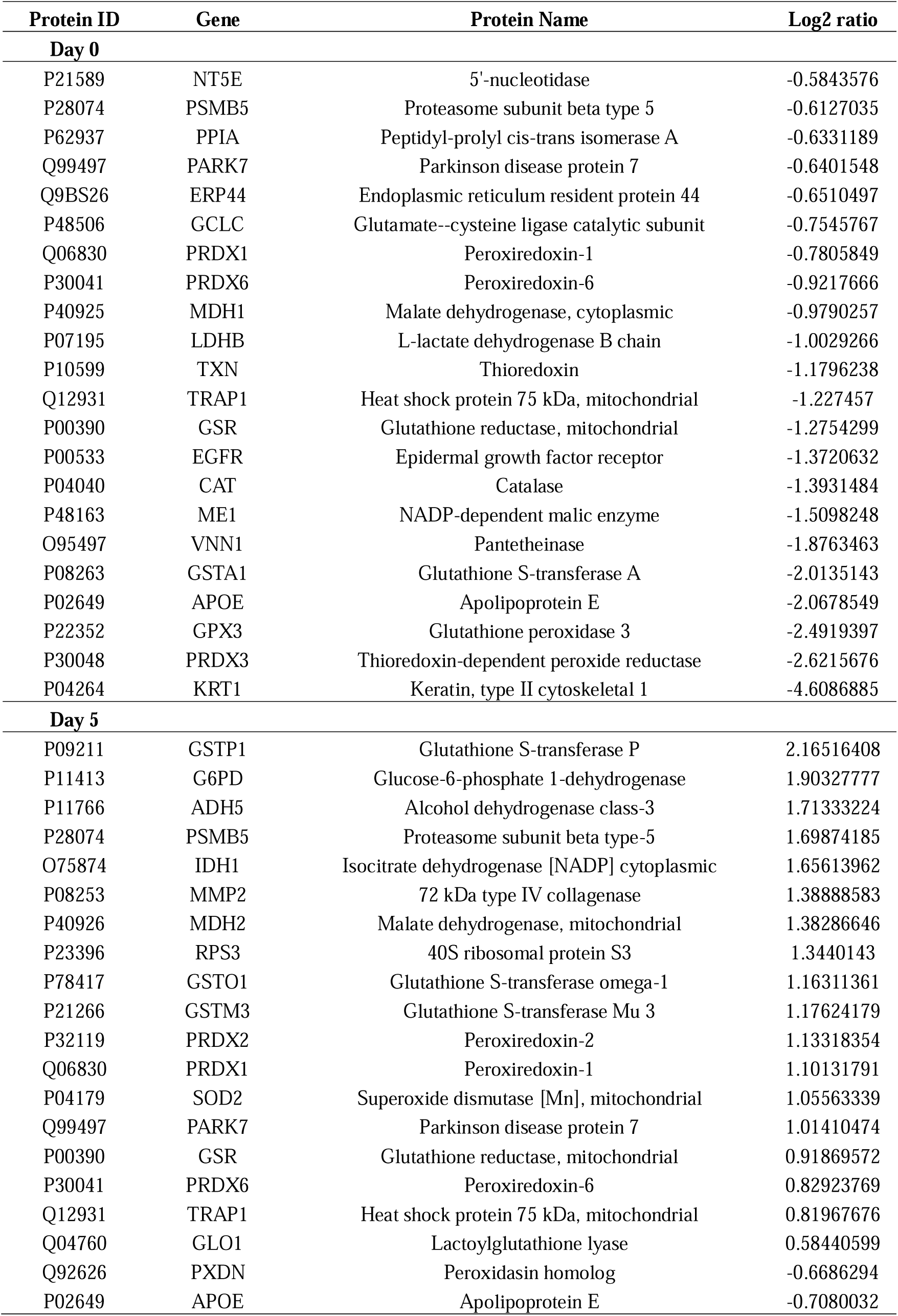
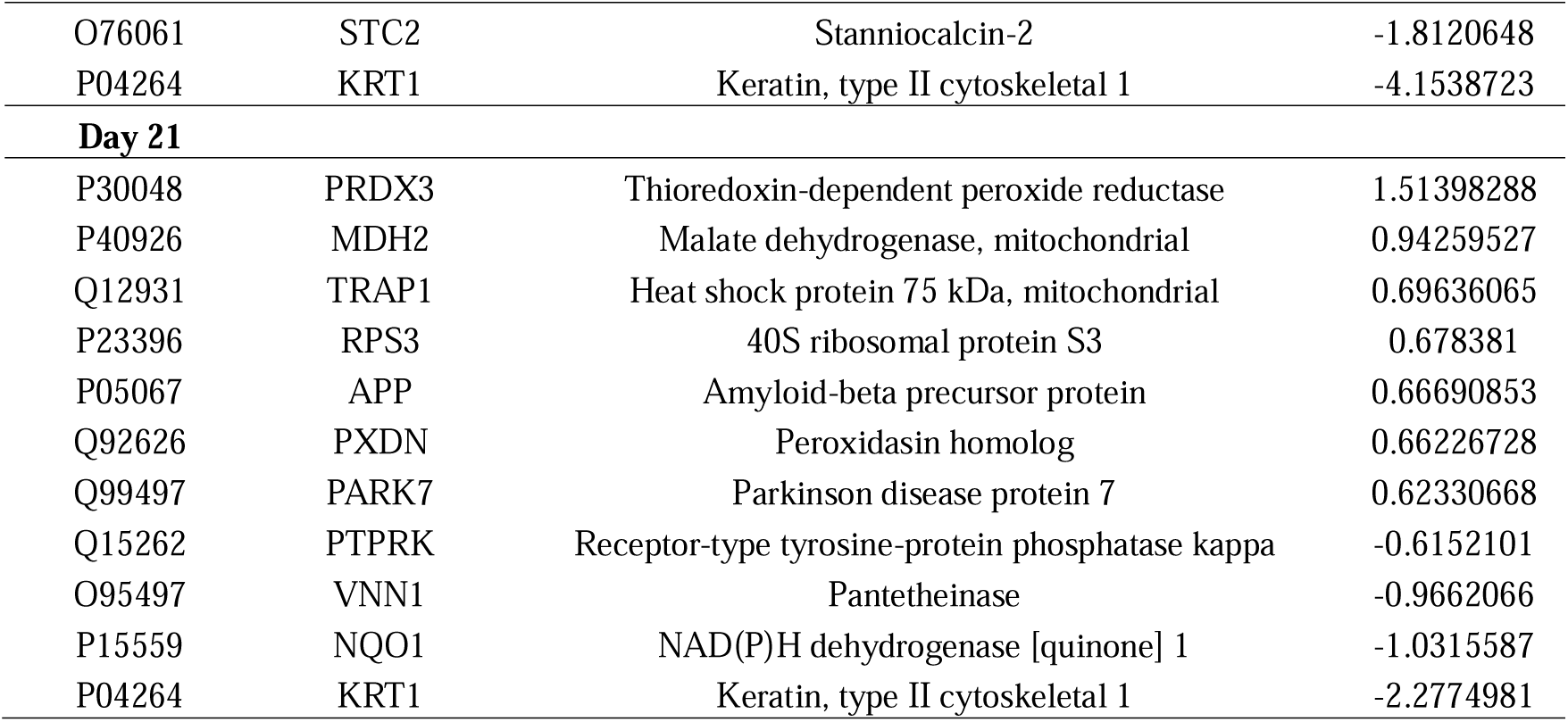
Upregulated and downregulated redox-related proteins in OB-MSCs versus NW-MSCs.

**Supplementary Table S5.**
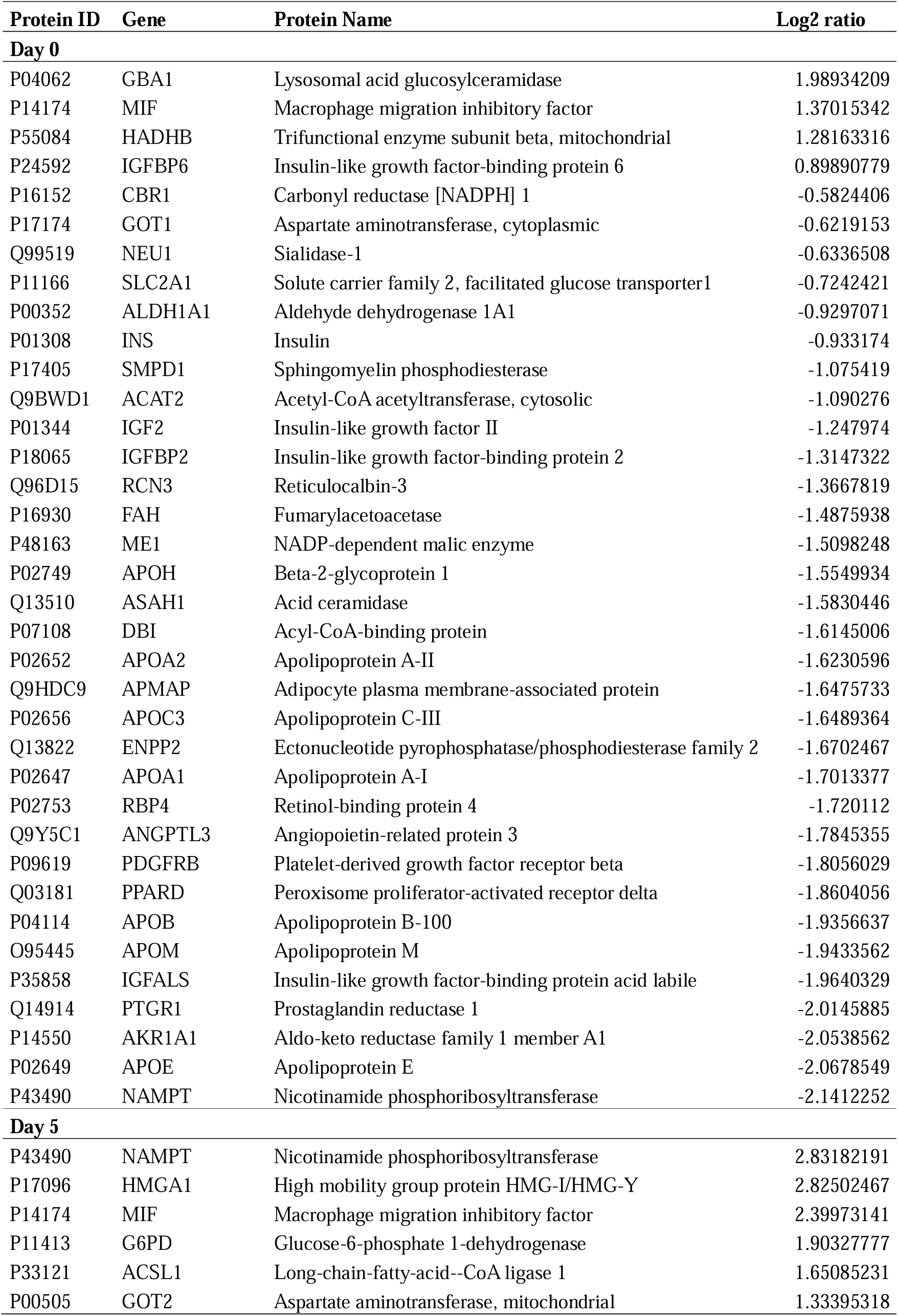
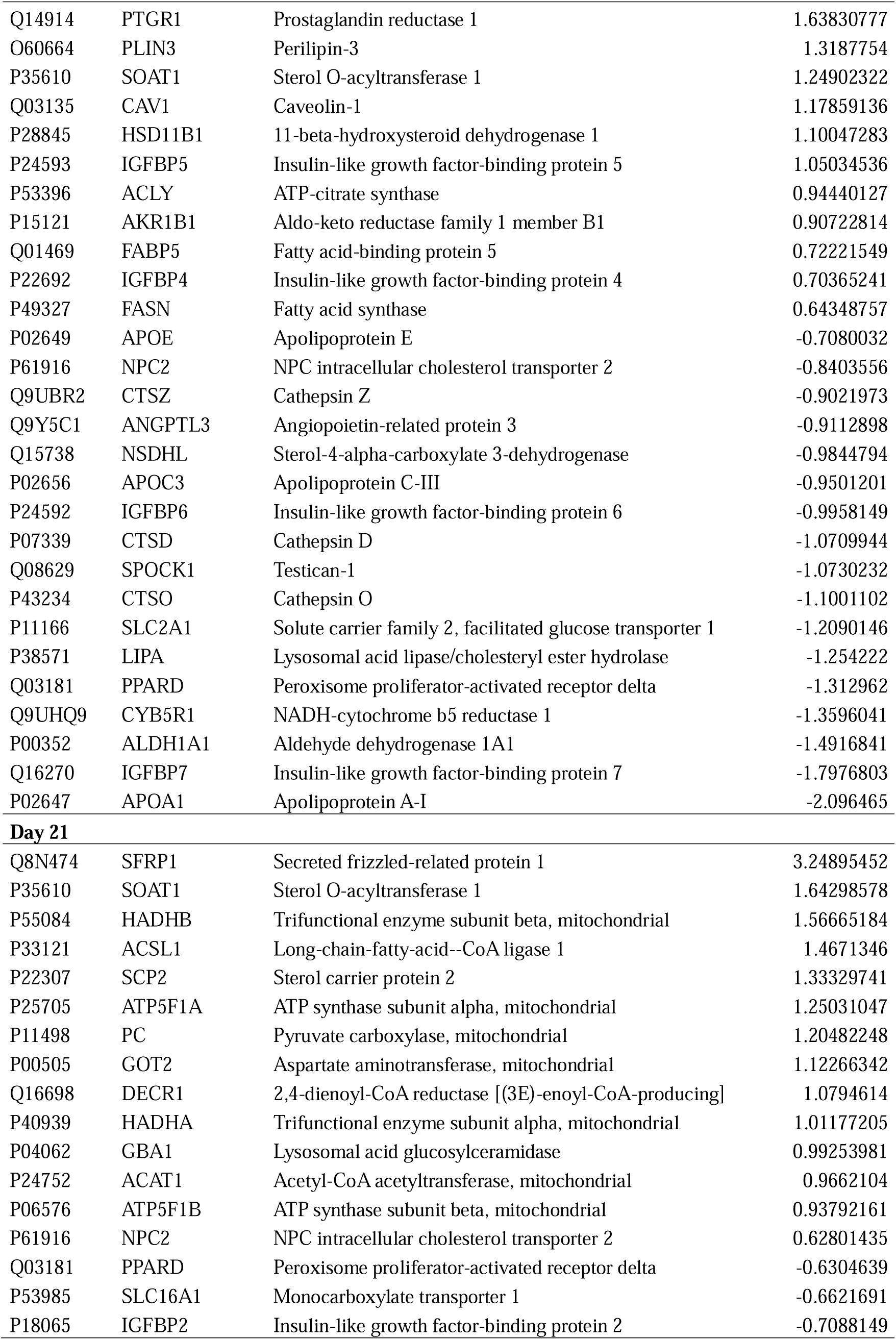
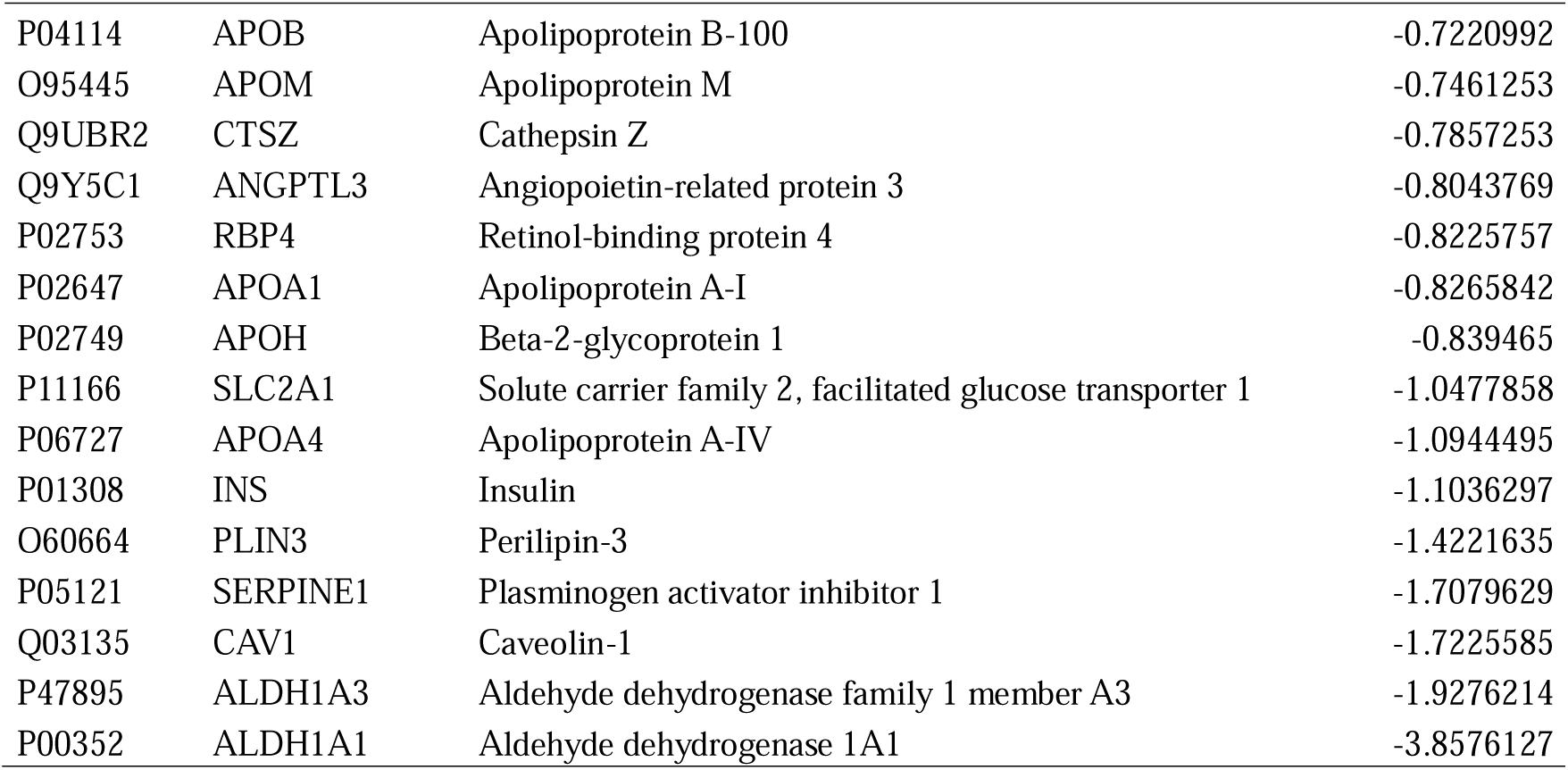
Upregulated and downregulated adipogenic-related proteins in OB-MSCs versus NW-MSCs.

