## Supplementary for "Profiling the secretome: maternal obesity impacts redox and adipogenic signaling molecules during neonatal mesenchymal stem cells’ adipogenesis"

### Supplementary Material

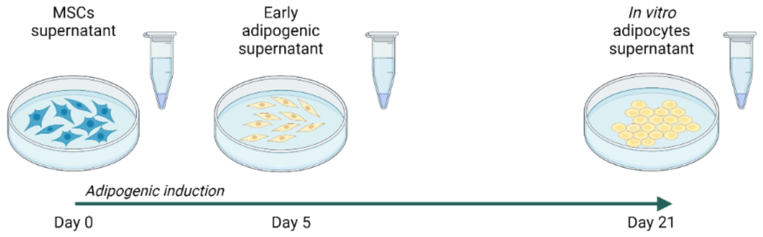


**Supplementary Figure S1.** Illustrative representation of the experimental design. NW-MSCs and OB-MSCs were induced for *in vitro* adipogenesis for 21 days. Supernatant was collected on day 0, 5 and 21 for secretomic analysis by LC-MS. Created in BioRender.com

**Supplementary Table S1. Top 20 subclass enrichment pathways of OB-MSCs versus NW-MSCs on day 0**

| GO | Category | Description | Count | Log10(P) | Log10(q) |
|---|---|---|---|---|---|
| R-HSA-6798695 | Reactome Gene Sets | Neutrophil degranulation | 61 | -36.06 | -31.80 |
| hsa04610 | KEGG Pathway | Complement and coagulation cascades | 32 | -34.88 | -30.92 |
| R-HSA-109582 | Reactome Gene Sets | Hemostasis | 65 | -33.25 | -29.48 |
| R-HSA-381426 | Reactome Gene Sets | Regulation of Insulin-like Growth Factor (IGF) transport and uptake by Insulin-like Growth Factor Binding Proteins (IGFBPs) | 33 | -30.31 | -26.65 |
| R-HSA-1474244 | Reactome Gene Sets | Extracellular matrix organization | 41 | -25.47 | -22.06 |
| R-HSA-422475 | Reactome Gene Sets | Axon guidance | 50 | -22.86 | -19.55 |
| GO:1900046 | GO Biol Processes | regulation of hemostasis | 21 | -20.62 | -17.60 |
| R-HSA-2173782 | Reactome Gene Sets | Binding and Uptake of Ligands by Scavenger Receptors | 17 | -19.52 | -16.66 |
| GO:0098609 | GO Biol Processes | cell-cell adhesion | 45 | -18.96 | -16.14 |
| hsa05322 | KEGG Pathway | Systemic lupus erythematosus | 25 | -18.83 | -16.02 |
| GO:1990748 | GO Biol Processes | cellular detoxification | 22 | -17.45 | -14.73 |
| hsa05171 | KEGG Pathway | Coronavirus disease - COVID-19 | 28 | -16.04 | -13.44 |
| R-HSA-140877 | Reactome Gene Sets | Formation of Fibrin Clot (Clotting Cascade) | 14 | -15.30 | -12.76 |
| GO:0019752 | GO Biol Processes | carboxylic acid metabolic process | 47 | -14.15 | -11.71 |
| GO:0043086 | GO Biol Processes | negative regulation of catalytic activity | 35 | -14.00 | -11.59 |
| R-HSA-3781865 | Reactome Gene Sets | Diseases of glycosylation | 21 | -13.87 | -11.47 |
| GO:0005975 | GO Biol Processes | carbohydrate metabolic process | 34 | -13.65 | -11.26 |
| R-HSA-5653656 | Reactome Gene Sets | Vesicle-mediated transport | 41 | -12.70 | -10.38 |
| GO:0030855 | GO Biol Processes | epithelial cell differentiation | 39 | -12.30 | -10.00 |
| R-HSA-196854 | Reactome Gene Sets | Metabolism of vitamins and cofactors | 22 | -12.03 | -9.76 |

**Supplementary Table S2. Top 20 subclass enrichment pathways of OB-MSCs versus NW-MSCs on day 5**

| GO | Category | Description | Count | Log10(P) | Log10(q) |
|---|---|---|---|---|---|
| R-HSA-422475 | Reactome Gene Sets | Axon guidance | 118 | -78.47 | -74.22 |
| R-HSA-6798695 | Reactome Gene Sets | Neutrophil degranulation | 88 | -51.84 | -48.90 |
| R-HSA-1474244 | Reactome Gene Sets | Extracellular matrix organization | 69 | -47.42 | -44.61 |
| R-HSA-381426 | Reactome Gene Sets | Regulation of Insulin-like Growth Factor (IGF) transport and uptake by Insulin-like Growth Factor Binding Proteins (IGFBPs) | 36 | -28.62 | -26.01 |
| R-HSA-195721 | Reactome Gene Sets | Signaling by WNT | 53 | -28.11 | -25.52 |
| R-HSA-5653656 | Reactome Gene Sets | Vesicle-mediated transport | 69 | -24.50 | -21.98 |
| R-HSA-109582 | Reactome Gene Sets | Hemostasis | 66 | -24.27 | -21.77 |
| GO:0097435 | GO Biol Processes | supramolecular fiber organization | 64 | -23.99 | -21.50 |
| R-HSA-71387 | Reactome Gene Sets | Metabolism of carbohydrates | 45 | -23.33 | -20.88 |
| GO:0005975 | GO Biol Processes | carbohydrate metabolic process | 54 | -23.11 | -20.69 |
| GO:0006457 | GO Biol Processes | protein folding | 39 | -22.38 | -19.97 |
| R-HSA-449147 | Reactome Gene Sets | Signaling by Interleukins | 52 | -19.86 | -17.52 |
| R-HSA-2173782 | Reactome Gene Sets | Binding and Uptake of Ligands by Scavenger Receptors | 17 | -16.72 | -14.46 |
| R-HSA-9711123 | Reactome Gene Sets | Cellular response to chemical stress | 32 | -16.20 | -13.97 |
| GO:0001568 | GO Biol Processes | blood vessel development | 48 | -15.08 | -12.93 |
| R-HSA-1280218 | Reactome Gene Sets | Adaptive Immune System | 59 | -15.02 | -12.88 |
| R-HSA-3000157 | Reactome Gene Sets | Laminin interactions | 14 | -14.94 | -12.80 |
| GO:0071826 | GO Biol Processes | protein-RNA complex organization | 30 | -13.10 | -11.09 |
| hsa04610 | KEGG Pathway | Complement and coagulation cascades | 19 | -13.07 | -11.06 |
| GO:0019752 | GO Biol Processes | carboxylic acid metabolic process | 56 | -12.84 | -10.84 |

**Supplementary Table S3. Top 20 subclass enrichment pathways of OB-MSCs versus NW-MSCs on day 21**

| GO | Category | Description | Count | Log10(P) | Log10(q) |
|---|---|---|---|---|---|
| R-HSA-156827 | Reactome Gene Sets | L13a-mediated translational silencing of Ceruloplasmin expression | 47 | -46.15 | -41.97 |
| R-HSA-1474244 | Reactome Gene Sets | Extracellular matrix organization | 65 | -42.87 | -39.32 |
| R-HSA-381426 | Reactome Gene Sets | Regulation of Insulin-like Growth Factor (IGF) transport and uptake by Insulin-like Growth Factor Binding Proteins (IGFBPs) | 41 | -35.09 | -32.02 |
| hsa05322 | KEGG Pathway | Systemic lupus erythematosus | 37 | -28.25 | -25.49 |
| R-HSA-109582 | Reactome Gene Sets | Hemostasis | 70 | -27.25 | -24.52 |
| R-HSA-6798695 | Reactome Gene Sets | Neutrophil degranulation | 61 | -26.61 | -23.88 |
| hsa04610 | KEGG Pathway | Complement and coagulation cascades | 27 | -22.71 | -20.09 |
| GO:0009611 | GO Biol Processes | response to wounding | 52 | -21.90 | -19.32 |
| R-HSA-3000178 | Reactome Gene Sets | ECM proteoglycans | 24 | -20.32 | -17.77 |
| GO:1900046 | GO Biol Processes | regulation of hemostasis | 22 | -18.48 | -15.97 |
| GO:0051604 | GO Biol Processes | protein maturation | 51 | -18.38 | -15.88 |
| GO:0030162 | GO Bioal Processes | regulation of proteolysis | 54 | -16.66 | -14.23 |
| GO:0019752 | GO Biol Processes | carboxylic acid metabolic process | 62 | -16.20 | -13.79 |
| GO:0001503 | GO Biol Processes | ossification | 36 | -15.79 | -13.43 |
| GO:0009725 | GO Biol Processes | response to hormone | 61 | -15.72 | -13.36 |
| R-HSA-2173782 | Reactome Gene Sets | Binding, Uptake of Ligands by Scavenger Recept | 16 | -15.25 | -12.93 |
| R-HSA-9694516 | Reactome Gene Sets | SARS-CoV-2 Infection | 36 | -15.12 | -12.81 |
| R-HSA-72649 | Reactome Gene Sets | Translation initiation complex formation | 18 | -15.10 | -12.79 |
| GO:1990748 | GO Biol Processes | cellular detoxification | 22 | -13.97 | -11.74 |
| hsa01200 | KEGG Pathway | Carbon metabolism | 21 | -12.63 | -10.49 |

**Supplementary Table S4. Upregulated and downregulated redox-related proteins in OB-MSCs versus NW-MSCs**

| Protein ID | Gene | Protein Name | Log2 ratio |
|---|---|---|---|
| **Day 0** | | | |
| P21589 | NT5E | 5'-nucleotidase | -0.5843576 |
| P28074 | PSMB5 | Proteasome subunit beta type 5 | -0.6127035 |
| P62937 | PPIA | Peptidyl-prolyl cis-trans isomerase A | -0.6331189 |
| Q99497 | PARK7 | Parkinson disease protein 7 | -0.6401548 |
| Q9BS26 | ERP44 | Endoplasmic reticulum resident protein 44 | -0.6510497 |
| P48506 | GCLC | Glutamate--cysteine ligase catalytic subunit | -0.7545767 |
| Q06830 | PRDX1 | Peroxiredoxin-1 | -0.7805849 |
| P30041 | PRDX6 | Peroxiredoxin-6 | -0.9217666 |
| P40925 | MDH1 | Malate dehydrogenase, cytoplasmic | -0.9790257 |
| P07195 | LDHB | L-lactate dehydrogenase B chain | -1.0029266 |
| P10599 | TXN | Thioredoxin | -1.1796238 |
| Q12931 | TRAP1 | Heat shock protein 75 kDa, mitochondrial | -1.227457 |
| P00390 | GSR | Glutathione reductase, mitochondrial | -1.2754299 |
| P00533 | EGFR | Epidermal growth factor receptor | -1.3720632 |
| P04040 | CAT | Catalase | -1.3931484 |
| P48163 | ME1 | NADP-dependent malic enzyme | -1.5098248 |
| O95497 | VNN1 | Pantetheinase | -1.8763463 |
| P08263 | GSTA1 | Glutathione S-transferase A | -2.0135143 |
| P02649 | APOE | Apolipoprotein E | -2.0678549 |
| P22352 | GPX3 | Glutathione peroxidase 3 | -2.4919397 |
| P30048 | PRDX3 | Thioredoxin-dependent peroxide reductase | -2.6215676 |
| P04264 | KRT1 | Keratin, type II cytoskeletal 1 | -4.6086885 |
| **Day 5** | | | |
| P09211 | GSTP1 | Glutathione S-transferase P | 2.16516408 |
| P11413 | G6PD | Glucose-6-phosphate 1-dehydrogenase | 1.90327777 |
| P11766 | ADH5 | Alcohol dehydrogenase class-3 | 1.71333224 |
| P28074 | PSMB5 | Proteasome subunit beta type-5 | 1.69874185 |
| O75874 | IDH1 | Isocitrate dehydrogenase [NADP] cytoplasmic | 1.65613962 |
| P08253 | MMP2 | 72 kDa type IV collagenase | 1.38888583 |
| P40926 | MDH2 | Malate dehydrogenase, mitochondrial | 1.38286646 |
| P23396 | RPS3 | 40S ribosomal protein S3 | 1.3440143 |
| P78417 | GSTO1 | Glutathione S-transferase omega-1 | 1.16311361 |
| P21266 | GSTM3 | Glutathione S-transferase Mu 3 | 1.17624179 |
| P32119 | PRDX2 | Peroxiredoxin-2 | 1.13318354 |
| Q06830 | PRDX1 | Peroxiredoxin-1 | 1.10131791 |
| P04179 | SOD2 | Superoxide dismutase [Mn], mitochondrial | 1.05563339 |
| Q99497 | PARK7 | Parkinson disease protein 7 | 1.01410474 |
| P00390 | GSR | Glutathione reductase, mitochondrial | 0.91869572 |
| P30041 | PRDX6 | Peroxiredoxin-6 | 0.82923769 |
| Q12931 | TRAP1 | Heat shock protein 75 kDa, mitochondrial | 0.81967676 |
| Q04760 | GLO1 | Lactoylglutathione lyase | 0.58440599 |
| Q92626 | PXDN | Peroxidasin homolog | -0.6686294 |
| P02649 | APOE | Apolipoprotein E | -0.7080032 |

**Supplementary Table S4.** Continued

| | | | |
|---|---|---|---|
| O76061 | STC2 | Stanniocalcin-2 | -1.8120648 |
| P04264 | KRT1 | Keratin, type II cytoskeletal 1 | -4.1538723 |
| **Day 21** | | | |
| P30048 | PRDX3 | Thioredoxin-dependent peroxide reductase | 1.51398288 |
| P40926 | MDH2 | Malate dehydrogenase, mitochondrial | 0.94259527 |
| Q12931 | TRAP1 | Heat shock protein 75 kDa, mitochondrial | 0.69636065 |
| P23396 | RPS3 | 40S ribosomal protein S3 | 0.678381 |
| P05067 | APP | Amyloid-beta precursor protein | 0.66690853 |
| Q92626 | PXDN | Peroxidasin homolog | 0.66226728 |
| Q99497 | PARK7 | Parkinson disease protein 7 | 0.62330668 |
| Q15262 | PTPRK | Receptor-type tyrosine-protein phosphatase kappa | -0.6152101 |
| O95497 | VNN1 | Pantetheinase | -0.9662066 |
| P15559 | NQO1 | NAD(P)H dehydrogenase [quinone] 1 | -1.0315587 |
| P04264 | KRT1 | Keratin, type II cytoskeletal 1 | -2.2774981 |

**Supplementary Table S5. Upregulated and downregulated adipogenic-related proteins in OB-MSCs versus NW-MSCs**

| Protein ID | Gene | Protein Name | Log2 ratio |
|---|---|---|---|
| **Day 0** | | | |
| P04062 | GBA1 | Lysosomal acid glucosylceramidase | 1.98934209 |
| P14174 | MIF | Macrophage migration inhibitory factor | 1.37015342 |
| P55084 | HADHB | Trifunctional enzyme subunit beta, mitochondrial | 1.28163316 |
| P24592 | IGFBP6 | Insulin-like growth factor-binding protein 6 | 0.89890779 |
| P16152 | CBR1 | Carbonyl reductase [NADPH] 1 | -0.5824406 |
| P17174 | GOT1 | Aspartate aminotransferase, cytoplasmic | -0.6219153 |
| Q99519 | NEU1 | Sialidase-1 | -0.6336508 |
| P11166 | SLC2A1 | Solute carrier family 2, facilitated glucose transporter1 | -0.7242421 |
| P00352 | ALDH1A1 | Aldehyde dehydrogenase 1A1 | -0.9297071 |
| P01308 | INS | Insulin | -0.933174 |
| P17405 | SMPD1 | Sphingomyelin phosphodiesterase | -1.075419 |
| Q9BWD1 | ACAT2 | Acetyl-CoA acetyltransferase, cytosolic | -1.090276 |
| P01344 | IGF2 | Insulin-like growth factor II | -1.247974 |
| P18065 | IGFBP2 | Insulin-like growth factor-binding protein 2 | -1.3147322 |
| Q96D15 | RCN3 | Reticulocalbin-3 | -1.3667819 |
| P16930 | FAH | Fumarylacetoacetase | -1.4875938 |
| P48163 | ME1 | NADP-dependent malic enzyme | -1.5098248 |
| P02749 | APOH | Beta-2-glycoprotein 1 | -1.5549934 |
| Q13510 | ASAH1 | Acid ceramidase | -1.5830446 |
| P07108 | DBI | Acyl-CoA-binding protein | -1.6145006 |
| P02652 | APOA2 | Apolipoprotein A-II | -1.6230596 |
| Q9HDC9 | APMAP | Adipocyte plasma membrane-associated protein | -1.6475733 |
| P02656 | APOC3 | Apolipoprotein C-III | -1.6489364 |
| Q13822 | ENPP2 | Ectonucleotide pyrophosphatase/phosphodiesterase family 2 | -1.6702467 |
| P02647 | APOA1 | Apolipoprotein A-I | -1.7013377 |
| P02753 | RBP4 | Retinol-binding protein 4 | -1.720112 |
| Q9Y5C1 | ANGPTL3 | Angiopoietin-related protein 3 | -1.7845355 |
| P09619 | PDGFRB | Platelet-derived growth factor receptor beta | -1.8056029 |
| Q03181 | PPARD | Peroxisome proliferator-activated receptor delta | -1.8604056 |
| P04114 | APOB | Apolipoprotein B-100 | -1.9356637 |
| O95445 | APOM | Apolipoprotein M | -1.9433562 |
| P35858 | IGFALS | Insulin-like growth factor-binding protein acid labile | -1.9640329 |
| Q14914 | PTGR1 | Prostaglandin reductase 1 | -2.0145885 |
| P14550 | AKR1A1 | Aldo-keto reductase family 1 member A1 | -2.0538562 |
| P02649 | APOE | Apolipoprotein E | -2.0678549 |
| P43490 | NAMPT | Nicotinamide phosphoribosyltransferase | -2.1412252 |
| **Day 5** | | | |
| P43490 | NAMPT | Nicotinamide phosphoribosyltransferase | 2.83182191 |
| P17096 | HMGA1 | High mobility group protein HMG-I/HMG-Y | 2.82502467 |
| P14174 | MIF | Macrophage migration inhibitory factor | 2.39973141 |
| P11413 | G6PD | Glucose-6-phosphate 1-dehydrogenase | 1.90327777 |
| P33121 | ACSL1 | Long-chain-fatty-acid--CoA ligase 1 | 1.65085231 |
| P00505 | GOT2 | Aspartate aminotransferase, mitochondrial | 1.33395318 |

**Supplementary Table S5.** Continued

| | | | |
|---|---|---|---|
| Q14914 | PTGR1 | Prostaglandin reductase 1 | 1.63830777 |
| O60664 | PLIN3 | Perilipin-3 | 1.3187754 |
| P35610 | SOAT1 | Sterol O-acyltransferase 1 | 1.24902322 |
| Q03135 | CAV1 | Caveolin-1 | 1.17859136 |
| P28845 | HSD11B1 | 11-beta-hydroxysteroid dehydrogenase 1 | 1.10047283 |
| P24593 | IGFBP5 | Insulin-like growth factor-binding protein 5 | 1.05034536 |
| P53396 | ACLY | ATP-citrate synthase | 0.94440127 |
| P15121 | AKR1B1 | Aldo-keto reductase family 1 member B1 | 0.90722814 |
| Q01469 | FABP5 | Fatty acid-binding protein 5 | 0.72221549 |
| P22692 | IGFBP4 | Insulin-like growth factor-binding protein 4 | 0.70365241 |
| P49327 | FASN | Fatty acid synthase | 0.64348757 |
| P02649 | APOE | Apolipoprotein E | -0.7080032 |
| P61916 | NPC2 | NPC intracellular cholesterol transporter 2 | -0.8403556 |
| Q9UBR2 | CTSZ | Cathepsin Z | -0.9021973 |
| Q9Y5C1 | ANGPTL3 | Angiopoietin-related protein 3 | -0.9112898 |
| Q15738 | NSDHL | Sterol-4-alpha-carboxylate 3-dehydrogenase | -0.9844794 |
| P02656 | APOC3 | Apolipoprotein C-III | -0.9501201 |
| P24592 | IGFBP6 | Insulin-like growth factor-binding protein 6 | -0.9958149 |
| P07339 | CTSD | Cathepsin D | -1.0709944 |
| Q08629 | SPOCK1 | Testican-1 | -1.0730232 |
| P43234 | CTSO | Cathepsin O | -1.1001102 |
| P11166 | SLC2A1 | Solute carrier family 2, facilitated glucose transporter 1 | -1.2090146 |
| P38571 | LIPA | Lysosomal acid lipase/cholesteryl ester hydrolase | -1.254222 |
| Q03181 | PPARD | Peroxisome proliferator-activated receptor delta | -1.312962 |
| Q9UHQ9 | CYB5R1 | NADH-cytochrome b5 reductase 1 | -1.3596041 |
| P00352 | ALDH1A1 | Aldehyde dehydrogenase 1A1 | -1.4916841 |
| Q16270 | IGFBP7 | Insulin-like growth factor-binding protein 7 | -1.7976803 |
| P02647 | APOA1 | Apolipoprotein A-I | -2.096465 |
| **Day 21** | | | |
| Q8N474 | SFRP1 | Secreted frizzled-related protein 1 | 3.24895452 |
| P35610 | SOAT1 | Sterol O-acyltransferase 1 | 1.64298578 |
| P55084 | HADHB | Trifunctional enzyme subunit beta, mitochondrial | 1.56665184 |
| P33121 | ACSL1 | Long-chain-fatty-acid--CoA ligase 1 | 1.4671346 |
| P22307 | SCP2 | Sterol carrier protein 2 | 1.33329741 |
| P25705 | ATP5F1A | ATP synthase subunit alpha, mitochondrial | 1.25031047 |
| P11498 | PC | Pyruvate carboxylase, mitochondrial | 1.20482248 |
| P00505 | GOT2 | Aspartate aminotransferase, mitochondrial | 1.12266342 |
| Q16698 | DECR1 | 2,4-dienoyl-CoA reductase [(3E)-enoyl-CoA-producing] | 1.0794614 |
| P40939 | HADHA | Trifunctional enzyme subunit alpha, mitochondrial | 1.01177205 |
| P04062 | GBA1 | Lysosomal acid glucosylceramidase | 0.99253981 |
| P24752 | ACAT1 | Acetyl-CoA acetyltransferase, mitochondrial | 0.9662104 |
| P06576 | ATP5F1B | ATP synthase subunit beta, mitochondrial | 0.93792161 |
| P61916 | NPC2 | NPC intracellular cholesterol transporter 2 | 0.62801435 |
| Q03181 | PPARD | Peroxisome proliferator-activated receptor delta | -0.6304639 |
| P53985 | SLC16A1 | Monocarboxylate transporter 1 | -0.6621691 |
| P18065 | IGFBP2 | Insulin-like growth factor-binding protein 2 | -0.7088149 |

**Supplementary Table S5.** Continued

| | | | |
|---|---|---|---|
| P04114 | APOB | Apolipoprotein B-100 | -0.7220992 |
| O95445 | APOM | Apolipoprotein M | -0.7461253 |
| Q9UBR2 | CTSZ | Cathepsin Z | -0.7857253 |
| Q9Y5C1 | ANGPTL3 | Angiopoietin-related protein 3 | -0.8043769 |
| P02753 | RBP4 | Retinol-binding protein 4 | -0.8225757 |
| P02647 | APOA1 | Apolipoprotein A-I | -0.8265842 |
| P02749 | APOH | Beta-2-glycoprotein 1 | -0.839465 |
| P11166 | SLC2A1 | Solute carrier family 2, facilitated glucose transporter 1 | -1.0477858 |
| P06727 | APOA4 | Apolipoprotein A-IV | -1.0944495 |
| P01308 | INS | Insulin | -1.1036297 |
| O60664 | PLIN3 | Perilipin-3 | -1.4221635 |
| P05121 | SERPINE1 | Plasminogen activator inhibitor 1 | -1.7079629 |
| Q03135 | CAV1 | Caveolin-1 | -1.7225585 |
| P47895 | ALDH1A3 | Aldehyde dehydrogenase family 1 member A3 | -1.9276214 |
| P00352 | ALDH1A1 | Aldehyde dehydrogenase 1A1 | -3.8576127 |
